# Presynaptic precursor vesicles originate from the *trans*-Golgi network, promoted by the small GTPase RAB2

**DOI:** 10.1101/2020.06.02.128991

**Authors:** Torsten W. B. Götz, Dmytro Puchkov, Janine Lützkendorf, Alexander G. Nikonenko, Christine Quentin, Martin Lehmann, Stephan J. Sigrist, Astrid G. Petzoldt

## Abstract

Reliable delivery of presynaptic material, including active zone and synaptic vesicle proteins from neuronal somata to synaptic terminals is prerequisite for faithful synaptogenesis and neurotransmission. However, molecular mechanisms controlling the somatic assembly of presynaptic precursors remain insufficiently understood. Here we show that in mutants of the small GTPase RAB2 active zone and synaptic vesicle proteins accumulated in the neuronal somata at the *trans*-Golgi network and were consequently depleted at synaptic terminals, provoking neurotransmission deficits. The ectopic presynaptic material accumulations consisted of heterogeneous vesicles and short tubules of 40×60 nm and segregated in subfractions either positive for active zone proteins or co-positive for synaptic vesicle proteins and LAMP1, a lysosomal membrane protein. Genetically, *rab2* behaved epistatic over *arl8*, a lysosomal adaptor controlling axonal export of precursors. Collectively, we here identified a Golgi-associated assembly sequence in presynaptic precursor vesicle biogenesis controlled by RAB2 dependent membrane remodelling and protein sorting at the *trans*-Golgi.

## Introduction

Synaptogenesis is based on the tightly controlled delivery of presynaptic material trafficking on presynaptic precursor vesicles (PVs) from their place of production, the neuronal soma, to the site of consumption, the synaptic terminal (Ahmari et al., 2000; Chia et al., 2013; Petzoldt et al., 2016; Sudhof, 2018). Impairments of presynapse formation and consequently presynaptic function cause severe neurodegenerative and neurodevelopmental diseases, such as Parkinson, ASD (Autism spectrum disorder), Alzheimer’s disease, mental retardation and micro- and macrocephaly (Parrini et al., 2016; Salinas et al., 2008; Waites and Garner, 2011). Different classes of presynaptic cargo proteins have to traffic coordinately along the axonal microtubules to the assembling presynapse (Maeder et al., 2014b) including active zone (AZ) scaffold proteins, large, multi-domain protein aiding the process of synaptic vesicle (SV) fusion, SV proteins, release factors and voltage gated calcium channels (VGCCs) (Fejtova and Gundelfinger, 2006; Petzoldt et al., 2016; Sudhof, 2012). Presynaptic proteins are thought to traffic on precursor vesicles in defined stoichiometric ratios as pre-assembled protein clusters associated with the vesicular membrane, providing building blocks for effective integration in the presynaptic specializations (Ahmari et al., 2000; Goldstein et al., 2008; Siebert et al., 2015; Tao-Cheng, 2007). Despite the fundamental importance of presynapse assembly, the cell-biological mechanisms underlying presynaptic precursor biogenesis are incompletely understood. Particularly debated are the cell biological origin, composition and cargo load of the precursors (Chia et al., 2013; Jin and Garner, 2008; Maeder et al., 2014a). To date, particularly *in vivo*, only sparse observations exist regarding presynaptic precursor assembly. Earlier studies had suggested that precursors may originate at the Golgi complex (Dresbach et al., 2006; Maas et al., 2012; Shapira et al., 2003). We recently found that presynaptic precursors possess a lysosomal membrane identity as they acquire lysosomal membrane proteins, but are devoid of the lysosomal peptidase cathepsin, in a hitherto unexplained maturation process, hence the term PLVs for presynaptic lysosome-related vesicles, before being axonally transported via the small Arf-like GTPase Arl8 (Vukoja et al., 2018). Arl8 is an evolutionary conserved adaptor for kinesin motor proteins implicated in the anterograde transport of lysosomes and lysosome-related organelles (Khatter et al., 2015; Rosa-Ferreira and Munro, 2011) and a similar Arl8-dependent mechanism regulates axonal SV precursor trafficking in *C. elegans* (Klassen et al., 2010).

Here we sought to identify novel regulators of precursor biosynthesis and hypothesized that as precursors are of vesicular nature their formation process might require small GTPases, such as RAB (“Ras-related in brain“) proteins. RAB proteins are crucial organizers of membrane trafficking between vesicular organelles by conferring identity to the vesicular structures they are bound to and controlling vesicle budding and fusion via effector protein recruitment such as sorting factors (Stenmark, 2009). RAB proteins function as molecular switches by changing from a GTP-bound active state to a GDP-bound inactive state and reverse, mediated by GTPase activating (GAP) proteins responsible for the GTP hydrolysis and guanine nucleotide exchange factors (GEFs) (Stenmark, 2009; Zhen and Stenmark, 2015).

Here we provide evidence that in an early step of the biosynthetic assembly pathway, presynaptic precursors originate from the *trans*-Golgi, requiring the highly conserved, Golgi-related GTPase RAB2 for assembly and maturation. RAB2 is reported as a Golgi resident and to act bidirectional in ER to Golgi trafficking (Liu and Storrie, 2012; Saraste, 2016; Tisdale and Balch, 1996), and in *Drosophila* RAB2 shows a neuronal expression pattern (Chan et al., 2011). Here we showed that presynaptic precursor formation requires a RAB2 dependent protein sorting and export process from the *trans*-Golgi network as an early step of precursor biogenesis. In conjunction with the previously shown function of RAB2 and RAB2 effectors in the promotion of dense core vesicle (DCV) maturation in neurons required for neuropeptide transport (Ailion et al., 2014; Edwards et al., 2009; Hannemann et al., 2012; Sumakovic et al., 2009) and the association of *rab2* mutations with neurodevelopmental defects, i.e. autism spectrum disorders (ASDs), schizophrenia (SCZ) (Kiral et al., 2018; Takata et al., 2016) and memory and prefrontal morphology defects (Li et al., 2015) in human, our data single out RAB2 as a crucial factor in early presynaptic precursor biogenesis.

## Results

Synaptic proteins, including both SV and AZ proteins are thought to transport from the somata along the axon to the presynapse on presynaptic precursor vesicles (PVs). To date, vesicular origin and molecular identity of these precursor vesicles are still debated. We sought to identify unknown molecular regulators of precursor biogenesis within the group of RAB proteins, small GTPases conferring identity, recruiting effectors and concerting various membrane trafficking events within cells (Zhen and Stenmark, 2015). More than 60 RAB proteins are described in human, while *Drosophila melanogaster* expresses 29 highly conserved RAB proteins (Jin et al., 2012). A common phenotype associated with the block of precursor vesicle transport, i.e. in the absence of kinesin motor protein UNC104/KIF1A (Zhangetal., 2017; Zhangetal., 2016) or the kinesin adaptor Arl8 (Vukoja et al., 2018), is the accumulation of presynaptic proteins in neuronal somata or proximal axons accompanied by a loss of these proteins from the respective synaptic terminals. We hence performed a motoneuron specific RNAi-knock-down screen and analyzed all neuronally expressed RAB proteins (Chan et al., 2011) for AZ and SV protein accumulations in motoneuron somata and associated depletions of these proteins at motoneuronal terminals of *Drosophila* 3^rd^ instar larvae. Of all RAB proteins tested (data not shown), exclusively RAB2 knockdown showed a clear combinatorial phenotype in the somata and at the synaptic terminals (see later in the text and Fig. S2A-F).

### Presynaptic proteins accumulate in the cell bodies of *rab2*^−/−^ deficient neurons

We thus analyzed a *rab2* null mutant created by open reading frame excision (Kohrs et al., bioRxiv), which we verified by Western Blot analysis (Fig. S1A), indeed proving absence of detectable RAB2 protein in this *rab2* mutant. The mutant animals were lethal at the early 3^rd^ larval stage in accordance with a previously described *rab2* null mutant (Lorincz et al., 2017). Immunofluorescence analysis of *rab2* mutant motoneuronal cell bodies in the peripheral cortex of larval ventral nerve cords (VNC) uncovered an ectopic accumulation of the AZ scaffold protein Bruchpilot (BRP) (Kittel et al., 2006; Wagh et al., 2006) (Fig. 1A). BRP did not accumulate diffusely in the cytoplasm of the neuronal cell body, but in distinct large aggregates not observed in controls (Fig. 1A, zoom). Both, BRP sum intensity, the integral of BRP mean pixel intensities over the VNC cortex (Fig. 1B) as well as BRP aggregate number (Fig. 1C) were strongly and significantly increased. We further analyzed the *rab2* mutant somata for a set of presynaptic proteins. The AZ scaffold protein RIM-binding protein (RIM-BP) (Acuna et al., 2015; Liu et al., 2011) accumulated in *rab2* mutant motoneuronal somata (Fig. 1D,E, S1B) and the large ectopic RIM-BP accumulations directly overlapped with the BRP signal (Fig. 1B). The essential AZ scaffold-associated release factor (m)UNC13A (Bohme et al., 2016) also strongly accumulated upon loss of RAB2 and UNC13A aggregates again overlapped with the BRP accumulations (Fig. S1G-I). SV proteins, here vesicular glutamate transporter protein (VGlut) (Santos et al., 2009), strongly accumulated in the *rab2*^−/−^ motoneuron somata as well (Fig. 1F, G, S1C). However, in contrast to RIM-BP and UNC13A, VGlut aggregates were 2-3-fold larger than BRP aggregates (Fig. S1D) and, interestingly, localized rather adjacent than overlapping relative to the BRP accumulations. Also other SV proteins, Synaptotagmin-1 (Syt-1), the SV release calcium sensor (Zhou et al., 2017) (Fig. S1J-L) and Dap160/intersectin (Fig. S1M-O), an endocytic component implied in SV recycling (Pechstein et al., 2010), accumulated in the neuronal somata of *rab2* mutants in a similar pattern as VGlut: in larger, but not directly overlapping aggregates adjacent to the BRP accumulations. Unlike SV and AZ proteins, levels of the endogenous mitochondrial ATP-Synthase, which also traffics from the cell body to the neuronal terminal, were not increased in *rab2* mutant motoneuronal somata (Fig. S1P-R). Thus, loss of RAB2 function specifically impairs presynaptic protein export, while other trafficking pathways are apparently not affected. We could previously show that axonally transported BRP positive precursors are co-positive for the lysosomal-associated membrane protein LAMP1 and the lysosomal marker Spinster, suggesting that presynaptic precursors entail membranes of lysosomal origin, hence their designation as PLVs (presynaptic lysosome-related vesicles) (Vukoja et al., 2018). Consequently, we here also tested for LAMP1 protein levels in *rab2* mutants by expression of an GFP-LAMP1 construct (Pulipparacharuvil et al., 2005) specifically in motoneurons. Compared to wild type brains, we observed a strong increase of the LAMP1 signal in the cell body of *rab2* mutants (Fig. 1H-I, S1E). Like SV proteins, LAMP1 positive aggregates were larger and localized adjacent to BRP accumulations. Our findings agree with the previous observations showing an increase of LAMP1 signal in VNCs of *rab2* mutant larvae (Lund et al., 2018). Notably, the ectopic presynaptic protein accumulations in *rab2* mutants were not positive for the standard marker of the lysosomal and autophagosomal machinery including the lysosomal peptidase CathepsinL (Fig. 1J,K, S1F), the adaptor protein of the autophagy-lysosomal pathway p62 (Fig. S1S-U) or ATG8a (Fig. S1V-X). None of these proteins were altered in their levels or localization upon loss of RAB2. Thus, the presynaptic protein accumulations we observe in *rab2^−/−^* deficient neurons do not represent degradative compartments but accumulations of LAMP1-positive presynaptic material.

**Fig. 1.**
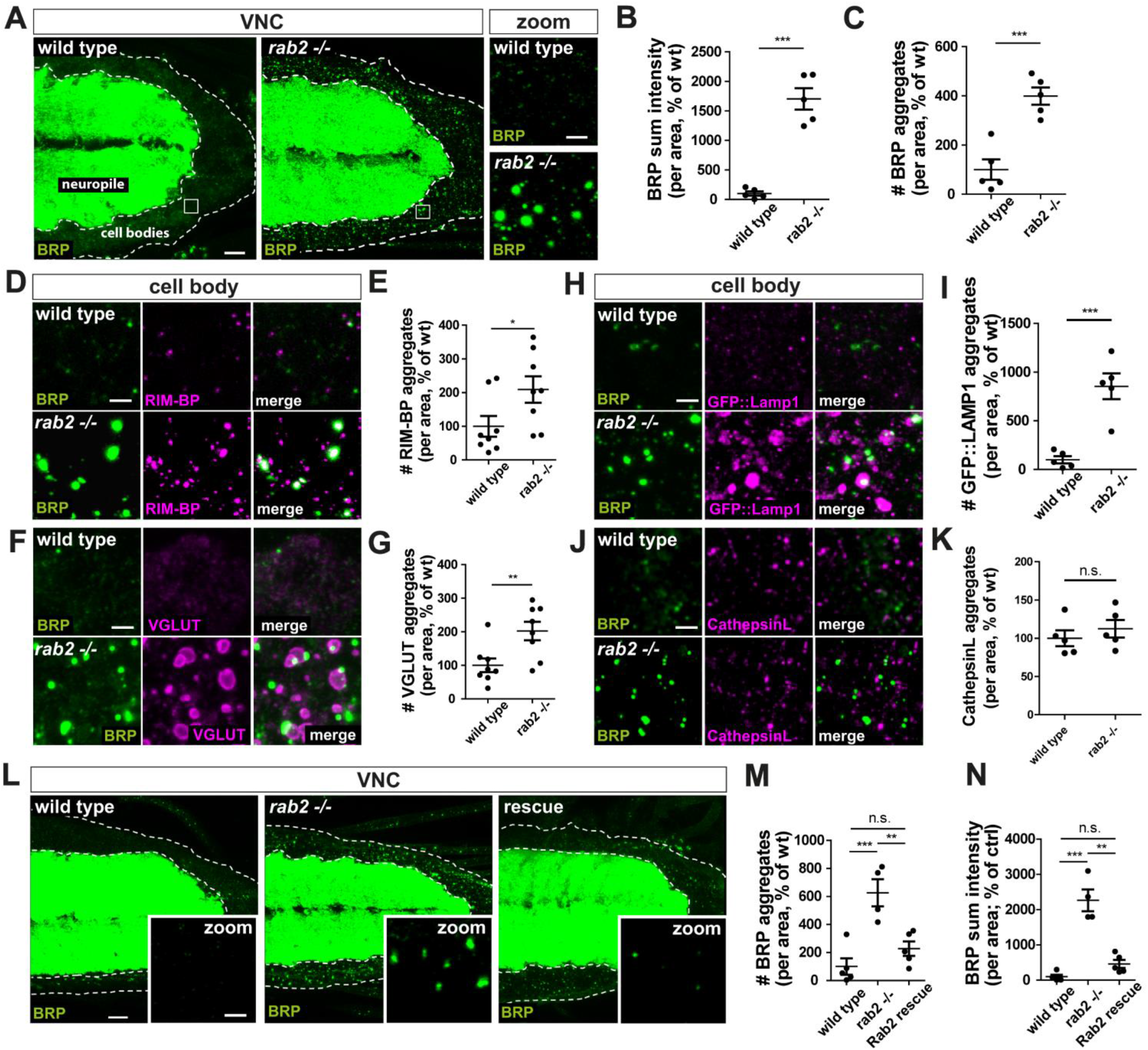
Presynaptic proteins accumulate in the cell bodies of *rab2*^−/−^ deficient neurons. (A-K) Accumulation of presynaptic proteins in motoneuronal somata of *rab2* mutant *Drosophila* larvae. (A) Confocalimages of larvalventralnervecord (VNC) from wild type and *rab2* mutant immunostained for the AZ scaffold protein BRP (green). BRP accumulated in large aggregates in the neuronalcellbodies localizedto the VNC cortex (dotted lines). White squares show zoom area. Scale bar, 10 μm, zoom 2 μm. (B,C) Quantifications of the representative images of (A). (B) Number of BRP aggregates (wt: 100.0 ± 40.93 %, n=5; rab2 −/− : 398.7 ± 35.27 %, n=5). (C) BRP sum intensity (wt: 100 ± 36.91 %, n=5; rab2 −/− : 1701 ± 181.7 %, n=5). (D) VNC immunostainings (zooms) co-labelling BRP (green) and the AZ scaffold protein RIM-BP (magenta) showed accumulation and co-localisation of RIM-BP-positive and BRP-positive areas. (E) Quantification of number of RIM-BP aggregates (wt: 100 ± 30.77 %, n=8; rab2 −/− : 209.1 ± 39.04 %, n=8). (F) VNC immunostainings co-labelling BRP(green) and the SV protein VGlut (magenta) or (H) lysosomal membrane protein GFP::LAMP1 (magenta) showed an accumulation of VGlut or LAMP1 in large aggregates adjacent to BRP aggregates. (G) Quantification of number of VGlut aggregates (wt: 100.0 ± 20.22 %, n=8; rab2 −/−: 202.2 ± 27.43 5 %, n=8) and (I) of LAMP1 aggregates (wt: 100.0 ± 36.72 %, n=5; rab2 −/−: 853.8 ±133.1 %, n=5). (J) VNC immunostainings co-labelling BRP (green) and the lysosomal peptidase CathepsinL (magenta) showed no increase or positional changes for lysosomes in *rab2* mutants. BRP aggregates of *rab2* mutants were not positive for CathepsinL. (K) Quantification of number of CathepsinL aggregates (wt: 100.0 ± 10.32 %, n=5; rab2 −/−: 112.4 ± 11.36 %, n=5). Scale bar D, F, H, J 2 μm. (L) Re-expression of endogenous levels of RAB2 rescued ectopic BRP accumulation in the cell bodies. Confocal images of the VNC from wild type, *rab2* mutant and *rab2* mutants with re-expressed RAB2 immunostained for BRP (green). Scale bar overview 10 μm, zoom 2 μm. (M,N) Quantifications of the representative images of (L). (M) Number of BRP aggregates (wt: 100.0 ± 58.90 %, n=5; rab2 −/−: 626.0 ± 96.90 %, n=4; Rab2 rescue: 227.4 ± 51.22 %, n=5). (N) BRP sum intensity (wt: 100.0 ± 53.28 %, n=5; rab2 −/−: 2262.0 ± 308.3 %, n=4; Rab2 rescue: 461.0 ± 114.2 %, n=5). All graphs show mean +/− SEM. Normality was tested with the Kolmogorov-Smirnov test, if normal distributed unpaired t-test (B, C, G, I, K; one-way ANOVA in M+N) was used, otherwise non-parametric Mann-Whitney test (E). *p<0.05, **p<0.01, ***p<0.001.

Importantly, we could “rescue” the BRP accumulation phenotype back to wild type BRP levels (Fig. L-N) by endogenous re-expression of RAB2 under the control of its own promotor in *rab2* mutants demonstrating the cell autonomous character of the *rab2* motoneuron phenotype. Finally, we used a cell specific RNA interference (RNAi) loss-of-function approach to directly and independently confirm the *rab2* mutant phenotype. Knock-down of RAB2 was verified by Western Blot analysis (Fig. S2A). We observed a significant accumulation of both AZ and SV proteins (BRP, VGlut and Syt-1) in the motoneuronal cell bodies (Fig. S2B-H), again indicative of a neuron specific, cell-autonomous RAB2 function. Collectively, we observed the accumulation of presynaptic proteins, including SV, AZ proteins, release factors and endocytic proteins in non-degradative aggregates in the neuronal somata upon loss of RAB2, a phenotype previously associated with defective presynaptic protein transport or assembly deficits of presynaptic precursors.

### Presynaptic biogenesis relies on the RAB2 dependent delivery of presynaptic material

If the accumulation of presynaptic proteins in the neuronal somata reflected deficits in regular assembly or transport of biosynthetic precursor material, we might expect a corresponding reduction of the respective proteins at the motoneuron synaptic terminals. We hence analyzed neuromuscular terminals of *rab2* mutant 3^rd^ instar larvae. Overall, the entire terminal of *rab2* mutants appeared thinner with atypically small presynaptic boutons, phenocopying larvae lacking kinesin Unc104/KIF1a (Pack-Chung et al., 2007; Zhang et al., 2016) or Arl8 (Vukoja et al., 2018). Strikingly, at *rab2* mutant synaptic terminals, overall BRP protein levels were reduced to only one third of wild type levels (Fig. 2A, Bi, C) and AZ numbers were strongly reduced (Fig. 2D). However, the size of the remaining AZ scaffolds was not altered (Fig. 2E), indicating that presynapse assembly per se was not affected by absence of RAB2, while the reduced availability of presynaptic biosynthetic material at the terminals accounted for defects in presynaptic biogenesis, evident in a reduced density of AZs forming. Apart from BRP, we also found other presynaptic proteins that accumulated in the cell bodies to be reduced at *rab2* mutant terminals: AZ scaffold protein RIM-BP (Fig. 2Bii,F,G), release factor UNC13A (Fig. S3A,E), SV proteins VGlut (Fig. 2Biii,H) and Syt-1 (Fig. S3B, F) as well as endocytic protein DAP160 (Fig. S3C,G). Importantly, mitochondrial ATP-Synthase, which did not accumulate in the cell bodies of *rab2* mutant animals (Fig. S1S-Q) was not decreased at the synaptic terminals (Fig. S3Di,ii,H), indicating that axonal transport is not generically affected by absence of RAB2.

**Fig. 2.**
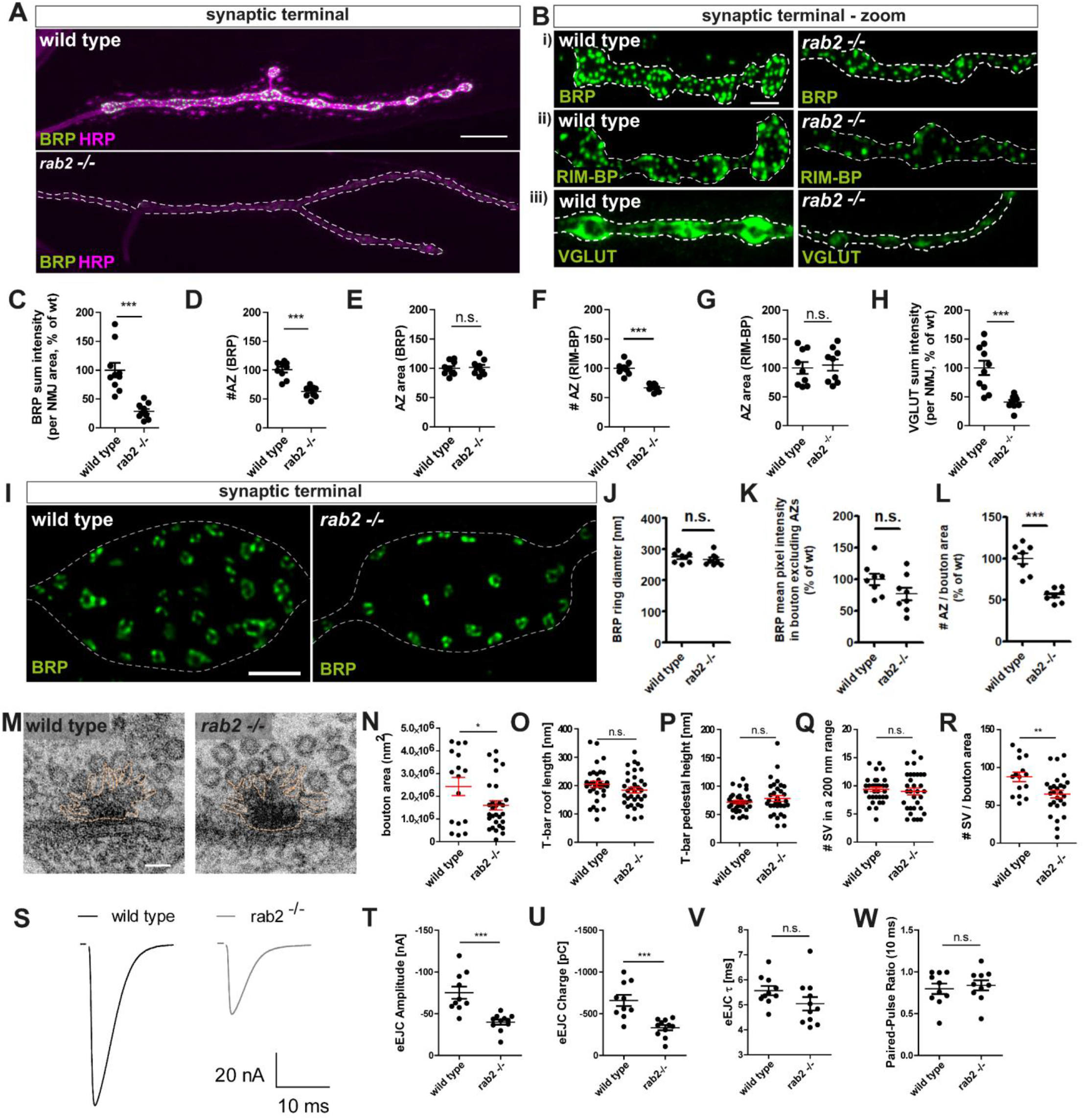
Presynaptic biogenesis relies on the RAB2 dependent delivery of presynaptic material. (A-G) Impaired presynaptic biogenesis in absence of RAB2 dependent presynaptic material delivery. (A, Bi) *Rab2* mutant NMJs (neuromuscular junctions) were thin and boutons small, showing reduced BRP level and AZ numbers. Confocal images of wild type and *rab2* mutant NMJs stained for BRP (green) and HRP (magenta), a neuronal membrane marker. Scale bar (A) 10 μm and (Bi) 3 μm. Also RIM-BP (green) (Bii) and VGlut (green) (Biii) levels are reduced at the NMJ. (C-E) Quantifications of the representative images of (A). (C) BRP sum intensity (wt: 100.0 ± 12.55 %, n=10; rab2 −/−: 28.61 ± 4.123 %, n=10). (D) Number of AZs labelled with BRP (wt: 100.8 ± 4.867 %, n=9; rab2 −/−: 63.06 ± 2.884 %, n=10). (E) AZ area (wt: 100.0 ± 3.552 %, n=10; rab2 −/−: 101.6 ± 4.314 %, n=9). (F) Quantification of the number of AZs labelled with RIM-BP (wt: 100.0 ± 4.051 %, n=8; rab2 −/−: 66.69 ± 2.279 %, n=8) and (G) AZ area (wt: 100.0 ± 10.29 %, n=9; rab2 −/−: 105.0 ± 9.855 %, n=9). (H) Quantification of VGlut sum intensity (wt: 100.0 ± 12.41 %, n=10; rab2 −/−: 40.82 ± 3.919 %, n=9). (I) Absence of RAB2 is not affecting AZ nano-architecture. Super-resolution STED microscopy images of wild type and *rab2* mutant NMJs labelled for BRP (green). Scale bar 1 μm. (J-L) Quantifications of the representative images of (I). (J) BRP ring diameter was not altered in *rab2* mutants (wt: 273.1 ± 5.671, n=8; rab2 −/−: 266.8 ± 6.697, n=8) and (K) neither did BRP accumulate outside of the AZs (wt: 100.0 ± 9.269, n=8; rab2 −/−: 77.36 ± 10.20, n=8), while number of AZs per bouton area was reduced (wt: 100.0 ± 6.286, n=8; rab2 −/−: 56.07 ± 2.833, n=8). (M-R) Ultrastructural analysis of wild type and *rab2* mutant presynapses showed an intact nano-architecture of the AZ scaffold and unaltered number of SVs. (N) Electron micrographs of wild type and *rab2* mutant AZs. (M-Q) Quantifications of the representative images of (M). (N) Reduced bouton area in *rab2* mutants (wt: 2.4×10^6^ ± 4×10^5^ nm^2^, n=16; rab2 −/−: 1.6×10^6^ ± 2×10^5^ nm^2^, n=29). (O) T-bar roof length (wt: 204.2 ± 10.65 nm, n=33; rab2 −/−: 183.7 ± 10.13 nm, n=34). (P) T-bar pedestal height (wt: 71.58 ± 3.036 nm, n=33; rab2 −/−: 78.24 ± 5.034 nm, n=34). (Q) Number of SVs within 200 nm of the T-bar centre (wt: 9.364 ± 0.3936, n=33; rab2 −/−: 9.000 ± 0.6140, n=32). (R) Number of SVs per bouton area (wt: 87.73 ± 6.303, n=15; rab2 −/−: 64.84 ± 5.076, n=27). (S-W) Impaired evoked neurotransmission due to absence of RAB2 measured by two electrode voltage-clamp electrophysiological recordings. (S) Representative example eEJC traces of wild type and *rab2* mutant synapses. (T) Reduced eEJC amplitudes of rab2 mutants (wt: −75.14 ± 7.416 nA, n=10; rab2 −/−: −39.82 ± 3.053 nA, n=11) and (U) reduced eEJC charge (wt: −657.7 ± 66.80 pC, n=10; rab2 −/−: −331.8 ± 31.78 pC, n=11). (V) Release kinetics with eEJC tau were not altered (wt: 5.567 ± 0.1846 ms, n=10; rab2 −/−: 5.042 ± 0.2695 ms, n=11), neither (W) short-term plasticity with a 10 ms paired-pulse ratio (wt: 0.7974 ± 0.06262, n=10; rab2 −/−: 0.8388 ± 0.05969, n=10). All graphs show mean +/− SEM. Normality was tested with the Kolmogorov-Smirnov test, if normal distributed unpaired two-tailed t-test (C-H, J-L, N-R, T-W) was used. *p<0.05, **p<0.01, ***p<0.001.

When we restricted RAB2 knock-down to the motoneuron via RNAi, we found BRP levels (Fig. S3I,J,K) and AZ number (Fig. S3L) similarly reduced as in the *rab2* mutant, but again AZ size was unaffected (Fig. S3M). Thus, RAB2 clearly operates in the delivery of presynaptic material in a cell autonomous manner.

To evaluate possible defects of the AZ architecture due to loss of RAB2 not detectable under standard optical resolution, we performed super-resolution STED (<u>st</u>imulated <u>e</u>mission <u>d</u>epletion) microscopy. The typical BRP C-terminal ring indicative for regular BRP incorporation and topology of the AZ scaffold (Fouquet et al., 2009; Kittel et al., 2006) appeared normal in *rab2* mutants (Fig. 2I) and BRP ring diameters were unchanged (Fig. 2J). Furthermore, we did not detect any increase in cytoplasmic BRP not integrated into AZs in NMJs boutons, which should be expected in case BRP cargo would reach the terminal but fail to appropriately incorporate in absence of RAB2 (Fig. 2K). In accordance with the confocal data, active zone densities were decreased at *rab2* mutant NMJ terminals (Fig. 2L).

To evaluate AZ architecture and SV distribution in detail, we performed ultrastructural analyses by electron microscopy (EM) of *rab2* mutant terminals (Fig. 2M). While bouton cross-sectional areas were reduced in *rab2* mutants (Fig. 2N), the architecture of the remaining AZs seemed not affected as the morphometric parameters of the electron dense T-bar, T-bar roof length and T-bar pedestal height were unaltered (Fig. 2O,P). Moreover, number of SVs physically attached at the T-bar were not changed (Fig. 2Q). Thus, consistent with the light microscopic data, ultrastructural analysis showed that the remaining presynapses assemble a normal AZ scaffold in RAB2 deficient neurons. However, the overall number of SVs per bouton area was significantly reduced (Fig. 2R), consistent with a reduction of delivered presynaptic SV-proteins (Fig. 2Biii, G). Taken together, data from all resolution levels collectively indicate that RAB2 is implied in the supply of biosynthetic presynaptic material to the synaptic terminal but clearly not in the regulation of “acute” protein incorporation at assembling presynapses.

We finally asked whether the remaining presynapses forming in *rab2* mutants would still be physiologically functional and performed two-electrode voltage clamp (TEVC) electrophysiological recordings to evaluate neurotransmission. Single action potential evoked response was reduced to half of the evoked excitatory junctional currents (aEJCs) in *rab2* mutants compared to wild type animals (Fig. 2S,T) and correspondingly also the evoked charge transfer (Fig. 2U). However, neither the eEJC kinetics (Fig. 2V) nor short term-plasticity (10 ms paired-pulse) (Fig. 2W) of release were altered in RAB2 deficient cells. RAB2-RNAi mediated knock-down specifically in neurons yielded similar results as the *rab2* null mutant: eEJC amplitude in knock-down animals was significantly reduced (Fig. S3N,O) as was the eEJC charge (Fig. S3P), while both kinetics (Fig. S3Q) and short term plasticity (Fig. S3R) of evoked release were unaltered. Thus, the reduction of available presynaptic material at the synaptic terminal due to absence of RAB2 clearly restricts neurotransmitter release, while those AZ still forming apparently remain functional, in accordance with the normal AZ architecture confirmed by microscopy.

Thus, absence of RAB2 provoked severe defects in presynaptic biogenesis due to shortage of presynaptic material at the terminal. An evident explanation for the combinatorial phenotype found in *rab2* mutants, that is accumulation of presynaptic proteins in the neuron cell bodies and simultaneous absence of the very same proteins from the synaptic terminal, was a function of RAB2 in the successful maturation of to be transported presynaptic precursors. To examine the relation of RAB2 and axonally transported precursor vesicles, we asked whether RAB2 as a membrane anchored GTPase might localize to *in vivo* trafficking precursors.

### RAB2 is a component of anterogradely trafficking presynaptic precursor vesicles

We first performed immunofluorescence analysis of *Drosophila* 3^rd^ instar larval axons, expressing RAB2 in a GTP locked, constitutively active form, tagged with a YFP (RAB2^QL^-YFP) in motoneurons and co-immunostained for BRP. Distinct RAB2^QL^ positive puncta were visible in the axon and we observed RAB2^QL^ and BRP co-positive puncta (Fig. 3A), also showed by line profile (Fig. 3B) and quantified with the Pearson’s correlation co-efficient (Fig. 3C). Similarly, we detected puncta co-positive for RAB2^QL^ and VGlut (Fig. 3D-F). We had previously shown that the lysosomal marker Spinster, (Rong et al., 2011; Sweeney and Davis, 2002) co-traffics anterogradely on PLVs (Vukoja et al., 2018). We hence expressed mRFP tagged Spinster in motoneurons and indeed detected RAB2^QL^ positive puncta co-localized with Spinster^mRFP^ (Fig. 3G-I) in axons. However, fixed samples intrinsically do not allow to determine, whether the observed organelles are mobile nor to analyze the directionality of movement. Thus, we decided to monitor axonal trafficking *in vivo* in intact living larvae (Andlauer and Sigrist, 2012a; Rasse et al., 2005). We generated animals co-expressing strawberry-tagged “BRP short” (BRP^D3^-straw in (Fouquet et al., 2009)) together with RAB2-YFP and confocally imaged axonal trafficking *in vivo*. We observed anterograde co-trafficking of BRP and RAB2 on vesicular organelles in the axon (Fig. 3J,K). Furthermore, RAB2 was also present on Spinster positive, anterogradely trafficking organelles (Fig. 3L,M). In summary, we suggest, that the RAB2 positive anterogradely trafficking organelles with a molecular signature of presynaptic transport vesicles and lysosome-related identity (according to the described PLVs) represent mature presynaptic precursor vesicles and will use this generic term accordingly from here on. Of note, we also observed retrograde co-positive BRP and RAB2 trafficking events (data not shown), which could reflect proteins cycling backwards from the terminal to the cell body for degradation, consistent with the conveyer belt hypothesis (Wong et al., 2012) or the back-transport of presynaptic material in degradative compartments (Jin et al., 2018).

**Fig. 3.**
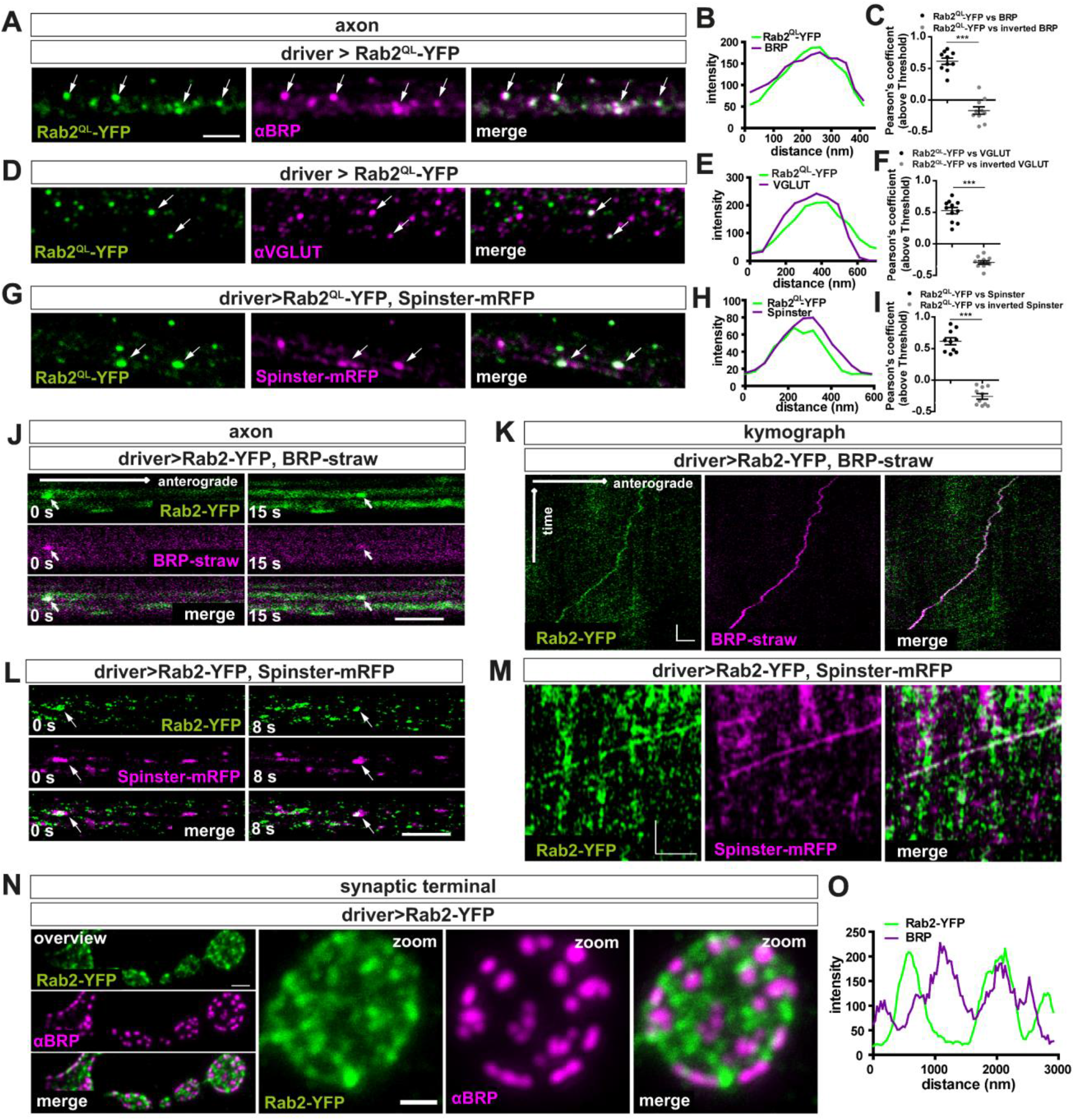
RAB2 is a component of anterogradely trafficking presynaptic precursor vesicles. (A-I) RAB2 co-localized with presynaptic proteins and lysosomal markers in the axon. Confocalimages showingpresynaptic transportvesicles in axons of 3 ^rd^ instar *Drosophila* larvae expressing RAB2^QL^-YFP. Immunofluorescent stainings showing several puncta (arrows) co-positive for RAB2^QL^-YFP (green) and (A) BRP (magenta), (D) VGlut or (G) the lysosomal marker Spinster. Scale bar 2 μm. (B,E,H) Corresponding line profiles for (A,D,G). (C,F,I) Corresponding Pearson’s correlation coefficients (C: Rab2^QL^-YFP vs. BRP: 0.62 ± 0.05, n=10; inverted BRP: −0.17 ± 0.06, n=10; F: Rab2^QL^-YFP vs. VGLUT: 0.53 ± 0.05, n=11; inverted VGLUT: −0.29 ± 0.03, n=11; I: Rab2^QL^-YFP vs. Spinster: 0.62 ± 0.06, n=10; inverted Spinster: −0.26 ± 0.04, n=10) confirming co-localization. (J-M) RAB2 co-trafficked on presynaptic transport vesicles positive for presynaptic proteins and lysosomal marker anterogradely along the axon *in vivo*. Confocal images from live imaged presynaptic precursors trafficking events (arrows) in larvae co-expressing RAB2-YFP (green) and (J) BRP^straw^ (magenta) and (L) Spinster^RFP^ (magenta). Stills at 0 s or 15 / 8 s respectively. Scale bar 3 μm. Kymograph of anterogradely trafficking PVs positive for RAB2-YFP (green) and (K) BRP^straw^ (magenta) or Spinster^RFP^ (magenta). Scale bar K: 3 μm / 10 sec; M: 4 μm / 20 sec. (N,O). RAB2 positive PVs reach the synaptic terminal. (N) Confocal images of 3 ^rd^ instar larval neuromuscular junctions (NMJs) expressing RAB2-YFP (green) stained for BRP (magenta). Scale bar overview 2 μm and zoom 1 μm. (O) Line profile of a corresponding synaptic bouton. Allgraphs show mean +/− SEM. Normality was tested with the Kolmogorov-Smirnov test and if normal distributed unpaired two-tailed t-test (C,F,I) was used. *p<0.05, **p<0.01, ***p<0.001.

If RAB2 traffics anterogradely on precursor vesicles, we should expect RAB2 to reach the synaptic terminal. Indeed, when we expressed RAB2-YFP into motoneurons we observed RAB2 in a punctate pattern at the synaptic terminals, interestingly localizing adjacent but not overlapping with the AZs (Fig. 3N,O). Potentially, RAB2 could designate here cargo loaded precursors waiting to be “discharged”.

In summary, we provided evidence that RAB2 as a membrane anchored protein associates with anterogradely trafficking presynaptic precursor vesicles, hence supporting the hypothesis that presynaptic protein accumulations in the somata of *rab2* mutants likely consist of precursors prematurely arrested in their assembly or biosynthetic maturation provoked by absence of RAB2.

### Presynaptic precursors are derived from the *trans*-Golgi network

To identify the molecular processes RAB2 is implied during its function in precursor formation, we first analyzed RAB2 distribution in motoneuron somata and expressed YFP-tagged RAB2 and detected the majority of RAB2 protein in large organelles in the cytoplasm (Fig. 4A, S4). To characterize the cell biological nature of these large organelles, we performed co-labeling analysis with common markers of the endo-lysosomal system, degradative organelles and the Golgi, using line profiles and quantifications of Pearson’s correlation coefficient for data validation. RAB2 did not co-localize with the early endosomal marker Rabenosyn-5 (Rbsn-5), a RAB5-effector protein, (Fig. S4A-C), the late endosomal marker Rab7 (Fig. S4D-F), the multivesicular body marker deep orange (DOR/VPS18)(Fig. S4G-I), the lysosomal marker CathepsinL (Fig. S4J-L) or the autophagosomal marker ATG8a (Fig. S4M-O). In contrast, we observed an explicit overlap of RAB2 with two Golgi markers, GM130 labeling the *cis*-(Fig. 4A-C) and Syntaxin16 (Syx16) the *trans*-Golgi (Fig.S4P-R). These findings agree with the previously described Golgi-related function of RAB2 (Liu and Storrie, 2012; Saraste, 2016; Tisdale and Balch, 1996). Of note, this does not, however, imply that smaller RAB2 positive signals or low abundance RAB2 protein might not localize to other cellular compartments as previously reported for other cell types, e.g. fat cells, nephrocytes, salivary glands or muscles (Ding et al., 2019; Fujita et al., 2017; Lorincz et al., 2017; Lund et al., 2018).

**Fig. 4.**
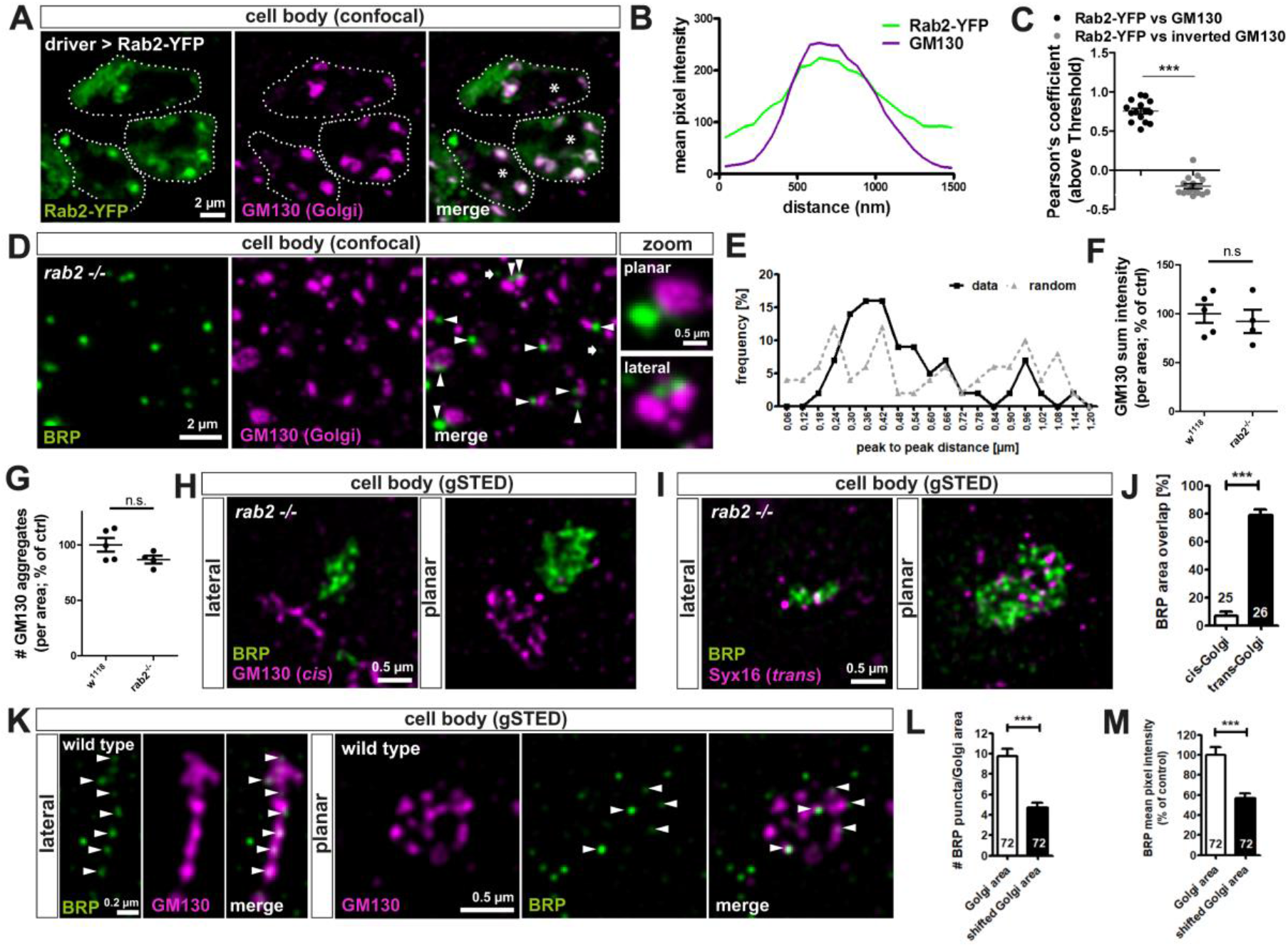
Presynaptic precursors are derived from the *trans*-Golgi network. (A-C) RAB2 localizes to the Golgi in motoneuronalcellsomata. (A) Confocalimages of neuronalcellbodies (outlined by dashed line, * nucleus) in the VNC cortex of 3 ^rd^ instar larvae expressing RAB2-YFP (green) stained for the Golgi marker GM130 (magenta) showing large RAB2 positive organelles co-positive for GM130. Scalebar 2μm. (B) Corresponding line profiles and (C) Pearson’s correlation coefficient (Rab2-YFP vs. GM130: 0.75 ± 0.04, n=15; inverted GM130: −0.21 ± 0.03, n=15) for (A) confirming localization of RAB2 to the Golgi. (D-E) Presynaptic precursor accumulations of *rab2* mutants localized adjacent to the Golgi. (D) Confocal images of neuronal cell bodies co-stained for BRP (green) and GM130 (magenta) with arrowheads highlighting adjacent localizations and arrows few non-adjacent events. Zoom shows planar and lateral orientation of the Golgi. Scale bar overview 2μm, zoom 0.5 μm. (E) Frequency of peak to peak distance distribution of BRP positive accumulations and Golgi f rom (D), black line for data and dashed grey line for 80 pixel shifted images (distance, % data (n), % shifted (n): 0.06 nm, 0.4%; 0.12 nm, 0.4%; 0.18 nm, 2.6%; 0.24 nm, 7.1%; 0.30 nm, 14.4%; 0.36 nm, 16.6%; 0.42 nm, 16.1%; 0.48 nm, 9.2%; 0.54 nm, 9.2%; 0.60 nm, 5.4%; 0.66 nm, 7.6%; 0.72 nm, 2.2%; 0.78 nm, 2.4%; 0.84 nm, 0.6%; 0.90 nm, 2.6%; 0.96 nm, 7.1%; 1.02 nm, 2.4%; 1.08 nm, 0.8%; 1.14 nm, 2.2%; 1.20 nm, 0.0%). N=84 single peak to peak measurements, data from 3 larvae, 2 VNC images each. (F,G) Quantification of Golgi parameters of *rab2* mutant brains from (D) showed no strong Golgi defects. (F) GM130 sum intensity (wt: 100.0 ± 9.326 %, n=5; rab2 −/−: 92.13 ± 11.92 %, n=4) and (G) G;130 aggregate number (wt: 100.0 ± 6.101 %, n=5; rab2 −/−: 86.70 ± 3.503 %, n=4) was not altered in *rab2* mutants. (H-J) BRP-positive precursors localized adjacent to the *cis*-Golgi and overlap with the *trans*-Golgi. STED images of *rab2* larval mutant brains co-stained for BRP (green) and (H) GM130 (*cis*-Golgi marker, magenta) and (I) Syx16 (*trans*-Golgi marker, magenta) in lateral or planar view of the Golgi. Scalebar 0.5μm. (J) Quantification of the area overlap of BRP with either *cis*- or *trans*-Golgi in % from (H,I) (*cis*-Golgi: 6.87 ± 2.82 %, n=25; *trans*-Golgi 79.08 ± 3.74 %, n= 26 single Golgi areas from 3 larvae and 2 VNC images each). (K-M) BRP positive puncta, possibly presynaptic precursors, were detectable in the cytoplasm of neuronal somata and enriched in the Golgi. (K) STED images of wild type larval brains co-stained for BRP (green) and GM130 (magenta) in lateral and planar view of the Golgi. Arrowheads pointon Golgi-localized PVs. Scalebar 0.5μm. (L) Quantification of with (K) number of PVs (Golgi area: 9.76 ± 0.71, shifted Golgi area: 4.74 ± 0.46, n=72 single Golgi areas from 3 larvae and 2 VNC images each) and (M) BRP mean pixel intensity within the Golgi-area compared to random shifted Golgi areas in % of control (Golgi area: 100.0 ± 7.62, shifted Golgi area: 56.72 ± 4.76, n=72 single Golgi areas from 3 larvae and 2 VNC images each). All graphs show mean +/− SEM. Normality was tested with the Kolmogorov-Smirnov test, if normal distributed unpaired two-tailed t-test (F+G) was used, otherwise non-parametric two-tailed Mann-Whitney test (C,J,L-M). *p<0.05, **p<0.01, ***p<0.001.

As RAB2 localizes to the Golgi, we consequently asked for the position of the BRP positive aggregates in relation to the Golgi in neuronal *rab2* deficient cells bodies, by co-labeling both BRP and Golgi. Importantly, we observed that BRP accumulations were not randomly distributed in the cytoplasm, but almost all BRP accumulations localized adjacent to the Golgi (Fig. 4D arrow heads, E), with a mean peak to peak distances between the *cis*-Golgi marker and BRP of typically around 400 nm (Fig. 4E). Thus, a Golgi-related function of RAB2 is clearly implied in the molecular processes causing the accumulation of presynaptic material at the Golgi.

RAB2 is described as a Golgi resident and its depletion in some cell types reported to affect Golgi apparatus integrity (Aizawa and Fukuda, 2015; Maringer et al., 2016). However, when comparing Golgi marker intensity and Golgi number of *rab2* mutants and wild type, no significant differences were detectable (Fig. 4F, G). Thus, from all we can tell, the general Golgi architecture was at least not severely affected in *rab2* mutants and presynaptic material accumulations do clearly not represent fragmented Golgi compartments.

Next, we wanted to distinguish whether the presynaptic biosynthetic material accumulated at the import (*cis*-) or export (*trans*-) side of the Golgi. Co-labeling of BRP with either *cis*- or *trans*-Golgi marker imaged by STED microscopy revealed that BRP positive accumulations were adjacent to the *cis*-Golgi marker GM130 both in lateral and planar views (Fig. 4H), but precisely overlapped with the *trans*-Golgi marker Syx16 (Fig. 4I), also validated by quantification (Fig. 4J). The *trans*-Golgi is the site of protein and lipid sorting for post-Golgi destinations, i.e. the plasma membrane, early, recycling and late endosomes, and export of different types of vesicles including secretory vesicles, lysosomal membrane protein (LMP) carriers and transport vesicles in a cell type specific manner (Keller and Simons, 1997; Klumperman, 2011; Pols et al., 2013). Thus, in an early step of precursor vesicle formation, presynaptic material could sort in a RAB2 dependent process from the *trans*-Golgi. If so, we should be able to detect at least a subfraction of presynaptic proteins at the Golgi of wild type neurons representing proteins being processed at or exported from the Golgi. Indeed, using STED microscopy, we detected BRP positive puncta in wild type cell bodies at the Golgi (Fig. 4K arrow heads), with an increased number and mean pixel intensity of BRP in the Golgi domain when compared to random shifted areas in the cytoplasm (Fig. 4L,M). Notably, however, we also observed BRP positive puncta in the cytoplasm (Fig. 4K, L). It is tempting to speculate that these cytoplasmic spots might represent post-Golgi precursors ready to be exported from the cell body.

In summary, our data so far provide evidence that RAB2 dependent membrane remodeling is required for presynaptic protein sorting from the *trans*-Golgi as an apparently early step of presynaptic precursor formation. Accumulating presynaptic material in *rab2* mutants likely represents immature precursor vesicles interrupted in their biosynthetic maturation process due to loss of the membrane trafficking regulator RAB2. However, the vesicular and membranous character of the presynaptic material accumulations in *rab2* mutants remain to be confirmed.

### Tubule-shaped vesicular membranes accumulate at the *trans*-Golgi in *rab2* mutants

We therefore turned towards ultrastructural analysis of wild type and *rab2* mutant neuronal cell bodies by electron microscopy. Interestingly, electron micrographs of *rab2* mutants showed a striking accumulation of circular to oval shaped vesicular membranes at the *trans*-Golgi (Fig. 5A,B, S5A,B), correlating with the position of presynaptic protein accumulations detected by light microscopy. In wild type cells, no vesicular accumulation at the Golgi of this kind were observed (Fig. 5A,B, S5A,B) also quantified in the significant increase in volume fraction of *trans*-Golgi network (TGN)-associated vesicles in the neuronal somata of *rab2* mutants (Fig. 5C). Together with the observation of axonally trafficking RAB2 positive precursor vesicles (Fig. 3J-M), we suggest that indeed somatic presynaptic material accumulations in RAB2 deficient neurons consist of immature precursor vesicles with a presynaptic cargo load. How do these maturation arrested, immature precursors of *rab2* mutants compare to the PLVs we previously described to accumulate in the cell body of *arl8* mutants (Vukoja et al., 2018)? The direct comparison of *arl8* mutant and *rab2* mutant electron micrographs revealed that PLVs detected in absence of Arl8 were with a ~60 nm short diameter significantly larger (Fig. 5B,D) than the ~40 nm sized *rab2* mutant precursors and possessed a more electron dense core (Fig. 5B) and a clearly more uniform, circular shape compared to the precursors detected in *rab2* mutants. The long diameters of both mutants were with ~75 nm almost identical (Fig. S5C).

**Fig. 5.**
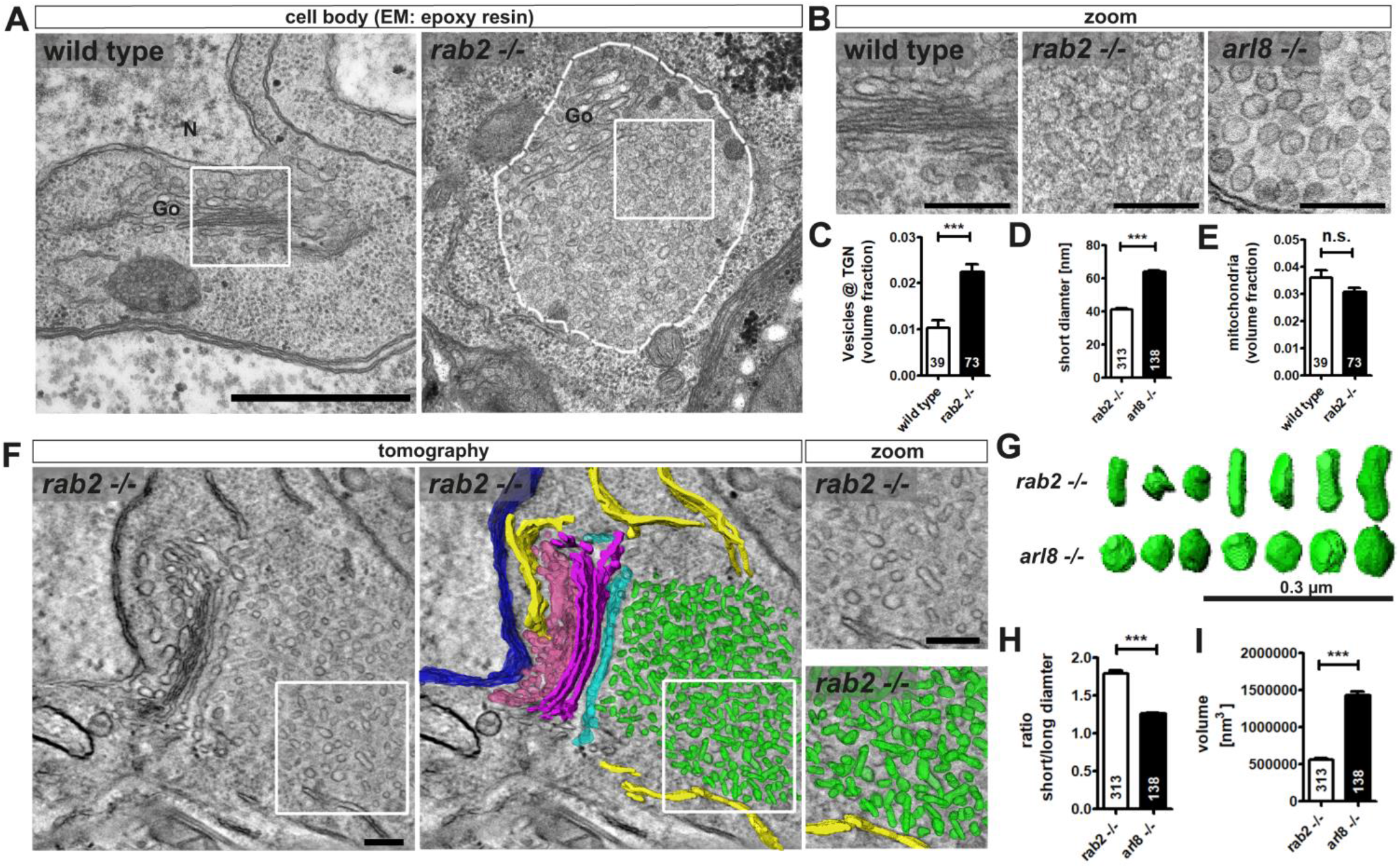
Tubule-shaped vesicular membranes accumulate at the *trans*-Golgi in *rab2* mutants. (A) Electron micrographs of wild type and *rab2* mutant cell bodies containing Golgi (Go) areas (N = nucleus) showed an accumulation of immature presynaptic precursors at the *trans*-Golgi (dashed line) in *rab2* deficient neurons. White squares indicate zoom area. Scale bar 500 nm. (B) Zoom shows no vesicle accumulations at the *trans*-Golgi in wild type somata (left). Biosynthetically arrested immature precursors of *rab2* mutants were clear cored with circular to oval shapes (middle) in contrast to mature PLVs of *arl8* mutants (right panel) which were larger, circular and partially with a dense core. Scale bar 100 nm. (C) Quantification of volume fraction of vesicles at the *trans*-Golgi network (TGN) showed an increase for *rab2* mutants (wild type: 0.0103 ± 0.0016, n= 39; rab2 −/−: 0.0224 ± 0.0018, n=73). (D) The short diameter of *trans*-Golgi area vesicles is shorter in *rab2* compared to *arl8* mutants (rab2−/−: 41.38 nm ± 0.51, n= 313; arl8−/−: 64.06 nm ± 0.8, n=138). N represents single vesicles. (E) Mitochondria volume fraction is not altered in *rab2* mutants compared to wild type (wild type: 0.036 ± 0.003, n= 39; rab2 −/−: 0.031 ± 0.002, n=73). (F) Example of TEM tomography reconstruction of the Golgi area of *rab2* mutants (left). Nucleus blue, ER yellow, ERGIC pink, *cis*-Golgi magenta, *trans*-Golgi cyan, early PVs green (right). White squares indicate zoom area. Scale bar overview and zoom 200 nm. Note the tubular structure of the immature precursors. Golgi architecture of *rab2* mutant resembled that of the wild type. (G) Comparison of representative random selection of 3D reconstructed *rab2* and *arl8* mutant precursors with an elongated, partially bulged shape for the former and circular vesicles for the latter. (H) Quantification of averaged *rab2* and *arl8* mutantvesicle short/longdiameter ratio (rab2−/−: 1.79 ± 0.03, n= 313; arl8−/−: 1.26 ± 0.02, n=138), n represents single vesicles, and (I) volume (rab2−/−: 0.56 *10^6^ nm^3^ ± 21040, n= 313; arl8−/−: 1.44 *10^6^ nm^3^ ± 48906, n=138), n represents single vesicles, accessed from 3D reconstructed vesicles confirm ellipticity and smaller volume of the immature precursors of *rab2* mutants compared to mature PLVs of *arl8* mutants. All graphs show mean +/− SEM. Normality was tested with the Kolmogorov-Smirnov test, if not normal distributed non-parametric two-tailed Mann-Whitney test was applied (C-E, H-I). *p<0.05, **p<0.01, ***p<0.001.

Notably, the volume fraction taken by mitochondria (Fig. 5E) was not altered in RAB2 deficient cells, confirming in agreement with the confocal data (Fig. S1E) that absence of RAB2 does not generically affect anterograde transport. However, we detected a 3fold increase in the incidence of somatic structures resembling autophagocytic organelles in *rab2* mutants (Fig. S5D), potentially interesting in connection with the previously reported function of RAB2 in autophagic degradative pathways (Ding et al., 2019; Fujita et al., 2017; Lorincz et al., 2017; Lund et al., 2018).

To deeper evaluate the apparent shape differences of precursor vesicles in *rab2* and *arl8* mutants we performed 3D electron tomography. Accumulating precursors in *rab2* mutants localized, in agreement with the STED light microscopic data (Fig. 4K,L), along the entire *trans*-Golgi, while the *cis*-Golgi facing the nucleus was devoid of such vesicular accumulations (Fig. 5F, S5E). Precursors were clearly detectable now in the 3D reconstruction as heterogeneous vesicular or short tubular structures (Fig. 5F zoom, 5G), with an average short/long diameter ratio of ~1.8 (Fig. 5G). In contrast, in *arl8* mutants, accumulating PLVs were rather homogenously shaped circular vesicles with short/long diameter ratio close to 1 (Fig. 5G,H). Vesicle volume quantifications showed that vesicles accumulating in *rab2* deficient cells were 3-fold smaller compared to those of *arl8* mutants (Fig. 5I) and had a 2-fold smaller surface area (Fig. S5F). Of note, the Golgi apparatus of *rab2* mutants did not appear fragmented and cisternae apparently were still forming normally (Fig. 5F), consistent with the confocal data (Fig. 4F,G).

In summary, we suggest that the tubule-shaped, clear cored vesicles of *rab2* mutants localized at the *trans*-Golgi with a smaller volume compared to the larger, strictly circular and dense-cored vesicles of *arl8* mutants represent immature and biosynthetically early arrested precursors, while *arl8* mutant vesicles constitute a more mature state in presynaptic precursor biogenesis. The immature precursors of *rab2* mutants could arise from a blockade of an early assembly step which requires a RAB2 catalyzed fission or fusion processes prior to later precursor maturation steps. We now utilized this RAB2 dependent premature arrest situation in biosynthetic precursor formation to assess in detail the molecular signature and cargo load of the retained precursors.

### Convergence of Golgi “trafficking streams” in the maturation of presynaptic precursors

Already the first standard confocal microscopy observations of accumulating presynaptic material in *rab2* mutants (see Fig. 1, S1) showed that some presynaptic cargoes were fully overlappingwith BRP, while others appeared to localize adjacent to the BRP signal, suggesting that variable Golgi-sorting routes might exist for different presynaptic proteins. To test this hypothesis, we first performed 3D confocal image reconstruction to investigate the presynaptic material distribution in the somatic accumulations of rab2 mutant VNCs in a three-dimensional representation. A 360° rotation of a 3D reconstructed precursor field co-stained for BRP and VGlut demonstrated that BRP positive and VGlut positive precursors occupied contiguous instead of overlapping areas (Fig. 6A), clearly visible in the line profile (Fig. 6B) and indicated by the low Pearson’s correlation co-efficient (Fig. 6C). Indeed, STED imaging further underlined this conclusion: BRP positive and VGlut positive precursors occupied neighboring areas (Fig. 6Di, Ei for line profile) and quantification of the area center distance of both precursor fields showed a clear shift of several hundred nanometers (Fig. 6J). In contrast, BRP and RIM-BP positive areas were precisely overlapping (Fig. 6Dii, Eii) and center distance between them nearly zero (Fig. 6J). Interestingly, LAMP1, expressed as GFP-LAMP1 construct in motoneurons, was not detectable in the BRP positive area (Fig. 6Diii, Eiii), but completely overlapped with the VGlut positive area (Fig. 6Div, Eiv), also evident in a large center distance between BRP and LAMP1 areas as compared to VGlut and LAMP1 distances (Fig. 6J). In a co-staining of LAMP1 with the Golgi marker GM130, LAMP1 positive areas consistently positioned at the Golgi of *rab2* mutants (Fig. S6A,B). Of note, a subfraction of the ectopic LAMP1 accumulations in *rab2* mutant were devoid of BRP signal and thus apparently not associated with early presynaptic precursor fields (Fig. S6C,D), potentially related to the previously described function of RAB2 in autophagic and lysosomal degradation (Ding et al., 2019; Fujita et al., 2017; Lorincz et al., 2017; Lund et al., 2018). In summary, these data suggest that at an early RAB2 dependent step of precursor biogenesis presynaptic proteins sort out of the Golgi on independent routes, separating active zone scaffold from SV/LAMP1 proteins.

**Fig. 6.**
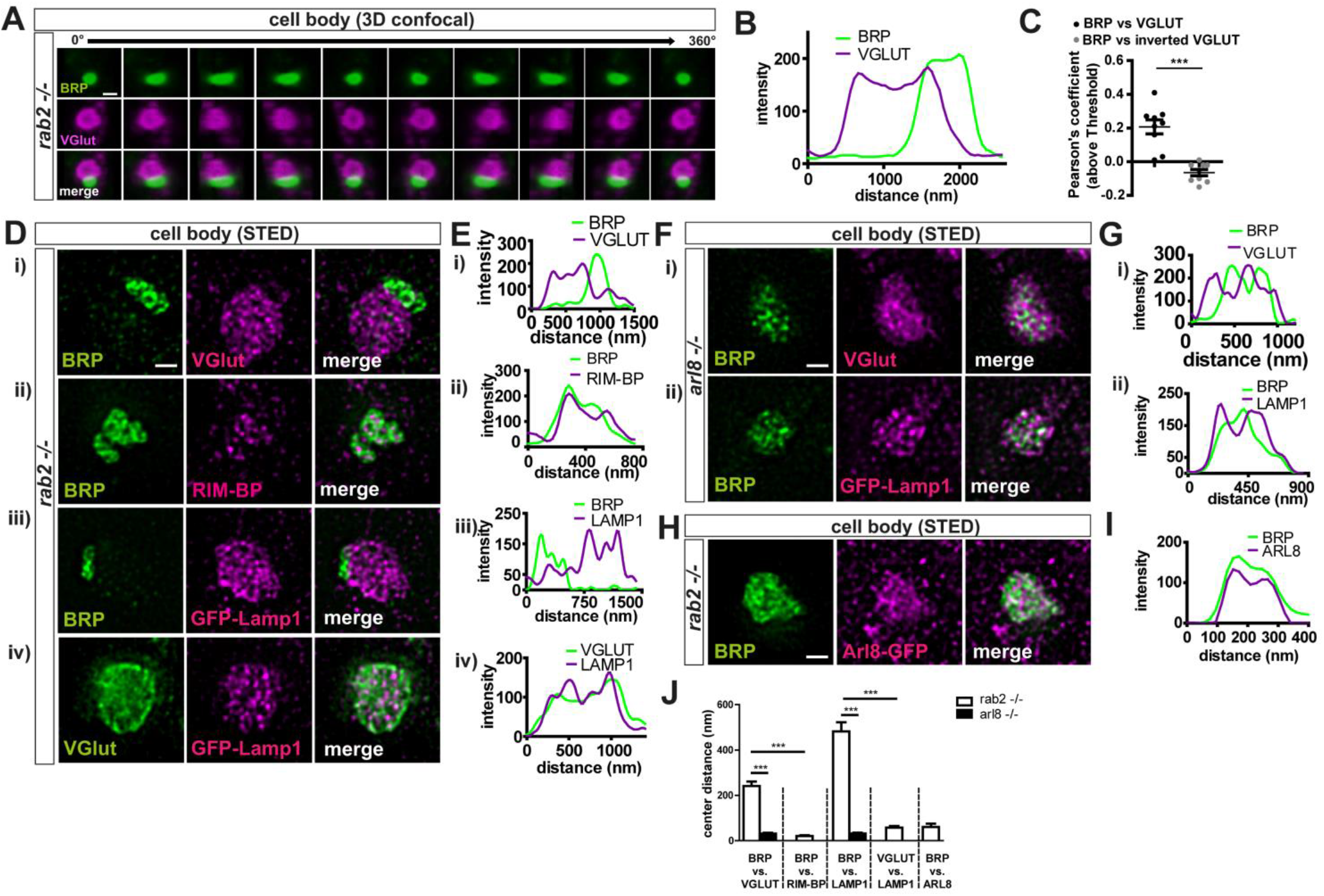
Convergence of Golgi “trafficking streams” in the maturation of presynaptic precursors. (A-C) BRP and VGlut cargo segregated in *rab2* mutant precursor accumulations. (A) 3D reconstruction of confocal images of a representative precursor accumulation in *rab2* mutant somata immunostained for BRP (green) and VGlut (magenta) in a 360° horizontal rotation showed two distinct either BRP or VGlut positive accumulation. Scale bar 1μm. (B) Corresponding line profiles and (C) Pearson’s correlation coefficients (BRP vs. VGlut: 0.21 ± 0.04, n=9; inverted VGlut: - −0.06 ± 0.02, n=9) for (A). (D,E) STED imaging of a representative precursor accumulation in *rab2* mutant somata immunostained for BRP (green) and (Di) VGlut (magenta), (Dii) RIM-BP (magenta), (Diii) LAMP1 (magenta, expressed as LAMP1-GFP construct) and (Div) VGlut (green) and GFP::LAMP1 (magenta). (Ei-iv) Corresponding line profiles. Of note, BRP and RIM-BP positive areas occupied overlapping areas, adjacent to VGlut and LAMP1 positive areas. (F-G) STED imaging of a representative precursor accumulation in *arl8* mutant somata immunostained for BRP (green) and (Fi) VGlut (magenta) and (Fii) GFP::LAMP1 (magenta). (Gi,ii) Corresponding line profiles. Mature PLVs had a uniform cargo load. (H,I) STED imaging of a representative immature precursor accumulation in *rab2* mutant somata immunostained for BRP (green) and Arl8-GFP (magenta, expressed as Arl8-GFP construct) and (I) corresponding line profile. Immature BRP-positive precursors of *rab2* mutants are Arl8-positive. Scalebar for (D,F,H) 0.5 μm. (J) Quantification of the area center distances of (D, F, G) (BRP vs. VGlut: rab2 −/−: 241.6 ± 19.00 nm, n=41; arl8 −/−: 30.85 ± 4.032 nm, n=34. BRP vs. RIM-BP: rab2−/−: 20.81 ± 2.951 nm, n=24. BRP vs. LAMP1: rab2 −/−: 482.0 ± 40.62 nm, n=28; arl8 −/−: 31.53 ± 4.20 nm, n=30. VGlut vs. LAMP1: rab2 −/−: 57.98 ± 6.93 nm, n=31. BRP vs. Arl8: rab2 −/−: 60.97 ± 14.16 nm, n=28). All graphs show mean +/− SEM. Normality was tested with the Kolmogorov-Smirnov test, if normal distributed unpaired t-test (C) was used, otherwise non-parametric Mann-Whitney test (J). *p<0.05, **p<0.01, ***p<0.001.

Interestingly, this segregation of presynaptic cargo distribution detected upon RAB2 depletion was not observed in the previously analyzed somata of *arl8* mutants (Vukoja et al., 2018). As this study was conducted with confocal microcopy, we also applied STED microscopy on *arl8* mutants for comparability. However, also upon a higher resolution, VGlut and LAMP1, again expressed as GFP-LAMP1 construct in the *arl8* mutant, nearly perfectly overlapped with BRP positive area in *arl8* mutants (Fig. 6Fi, ii, Gi, ii) with a small center distances (Fig. 6J), clearly contrasting *rab2* mutants. Finally, the question arose whether the immature presynaptic precursors of *rab2* mutants were already charged with the kinesin adaptor Arl8 or would gain the mobility factor at a later biosynthetic maturation step. We expressed a GFP tagged Arl8 (Vukoja et al., 2018) in *rab2* mutant animals and found Arl8 already present on immature precursors (Fig. 5H,I, J), restricted to the BRP positive area, hence a component of the subfraction of precursors charged with AZ scaffold proteins.

Collectively, these data suggest that precursor vesicle proteins in a first biosynthesis step sort from the *trans*-Golgi on at least two distinct routes, a process regulated by the small GTPase RAB2, as in absence of RAB2, immature, small and tubule-shaped precursors with distinct cargo accumulate. In subsequent precursor vesicle maturation steps, immature precursors converge to mature, larger, circular and Arl8-dependently transportable PLVs with a uniform presynaptic protein cargo load and a lysosomal membrane identity, possibly by direct fusion or through conventional fusion to the end/lysosomal compartment and subsequent segregation. Consequently, this would suggest that RAB2 functions upstream of Arl8 during precursor biogenesis. We hence performed genetic epistasis experiments to test this hypothesis.

### RAB2 acts upstream of Arl8 in presynaptic precursor biogenesis

We created a *rab2/arl8* double mutant and compared the cell body presynaptic protein accumulation phenotype to the respective single mutants. The double mutant showed a higher lethality than the single mutants (*rab2* mutant larvae did survive till to the early 3^rd^ instar larval stage, while *arl8* mutant larvae died earlier as 2^nd^ instar larvae) with only very few larvae surviving until the late 2^nd^ instar larval stage. In the cell bodies we observed that both number of BRP accumulations and BRP sum intensity were 2-3 times increased in *arl8* compared to *rab2* mutants (Fig. 7A-C). We used this phenotypic difference to distinguish between a *rab2*- and an *arl8*-related phenotype. Notably, the *rab2/arl8* double mutant larvae clearly resembled the single *rab2* mutant and we detected no significant difference between the single *rab2* and the *rab2/arl8* double mutant regarding both BRP sum intensity and AZ number in the VNC (Fig. 7A-C)). Thus, the interruption of precursor biogenesis due to absence of RAB2 prevents the entrance in the later Arl8 dependent biosynthetic maturation steps, placing RAB2 function upstream of Arl8 during presynaptic precursor biogenesis. Consequently, when analyzing the synaptic terminals of *rab2, arl8* double mutants, no additive effect in the reduction of the synaptic material was observed (Fig. 7D,E), again consistent with our interpretation that indeed both proteins act in a sequential rather than in parallel.

**Fig. 7.**
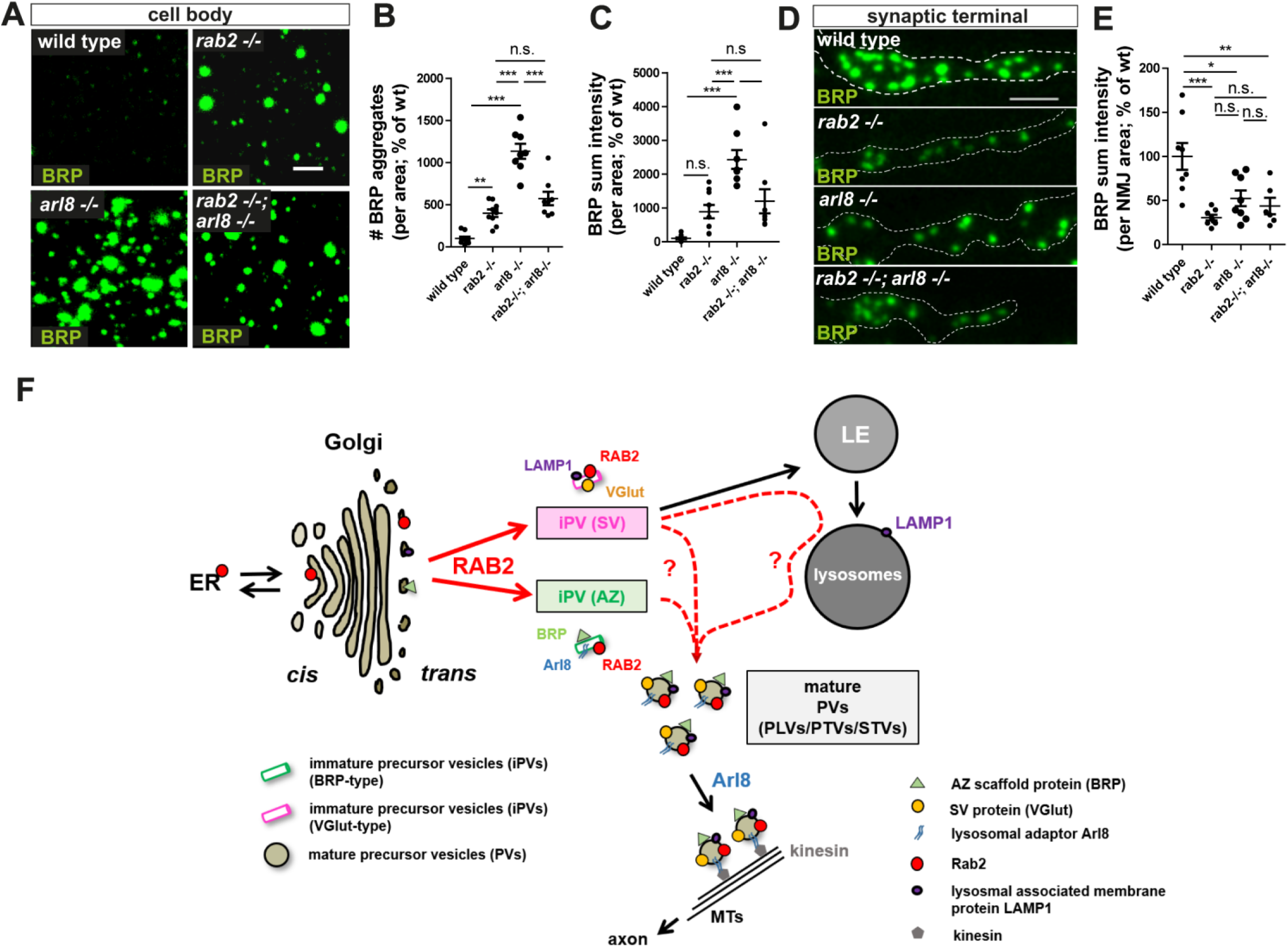
RAB2 acts upstream of Arl8 in presynaptic precursor biogenesis. (A-E) Genetic epistasis experiment of RAB2 and Arl8 comparing wild type, single *rab2* and *arl8* mutants with *rab2/arl8* double mutant larvae for cell body and synaptic terminal phenotypes. (A) Confocal images of the neuronal somata stained for BRP (green). Scalebar 2 μm. (B) Quantification of the number of BRP aggregates (wt: 100 ± 23.17 %, n=9; rab2 −/−: 399.3 ± 44.69 %, n=9; arl8 −/−: 1134 ± 89.10 %, n=8; rab2 −/−, arl8 −/− : 573.9 ± 79.89 %, n=8) and (C) BRP sum intensity (wt: 100 ± 32.66 %, n=9; rab2 −/−: 889.8 ± 198.3 %, n=9; arl8 −/−: 2435 ± 276.1 %, n=8; rab2 −/−, arl8 −/− : 1200 ± 355.5 %, n=8) sown in (A). The *rab2/arl8* double mutant reflects the single *rab2* mutant phenotype, placing RAB2 function upstream of Arl8. (D) Confocal images of the synaptic terminal stained for BRP (green). Scalebar 3 μm. (E) Quantification of the BRP sum intensity (wt: 100 ± 15.22 %, n=8; rab2 −/−: 30.56 ± 3.18 %, n=8; arl8 −/−: 52.43 ± 8.86 %, n=8; rab2 −/−, arl8 −/− : 43.74 ± 9.36 %, n=8) of (D). Both single mutants and the *rab2/arl8* double mutant show an equal reduction of BRP levels, indicating that RAB2 and Arl8 act sequentially during presynaptic precursor biogenesis. (F) Model of presynaptic precursor biogenesis. All graphs show mean +/− SEM. Normality was tested with the Kolmogorov-Smirnov test, if normal distributed one-way ANOVA was used (B+C). *p<0.05, **p<0.01, ***p<0.001.

## Discussion

Presynaptic proteins need to be transported on presynaptic precursor vesicles from the neuronal soma to the site of consumption, the synaptic terminal. And, although of fundamental importance, remained our understanding of the cell biological origin and molecular mechanics of presynaptic precursor biogenesis fragmentary. We here provide evidence, that in an early step of precursor biogenesis synaptic proteins sort out of the secretory *trans*-Golgi through independent routes on early,small and tubular membranous vesicles, a process clearly regulated by the small Golgi-related and highly conserved GTPase RAB2 (Chan et al., 2011; Lardong et al., 2015; Liu and Storrie, 2012; Lorincz et al., 2017; Saraste, 2016; Tisdale and Balch, 1996). Subsequent fusion steps of the post-Golgi early precursors could yield either directly by mutual fusion or via the endo-lysosomal compartments, the mature, full sized and uniformly charged presynaptic precursor vesicles (Fig. 7F).

Several independent lines of evidence, including confocal, super-resolution light and electron microscopy, live imaging in *Drosophila* larvae and genetic assays, support this model: biosynthetic presynaptic material including AZ scaffold, SV proteins, release factors and endocytic presynaptic proteins, accumulated ectopically in the neuronal cell body in absence of RAB2 (Fig. 1A-G, S1A-O). These presynaptic accumulations do clearly not represent degradative compartments as they were devoid of lysosomal and autophagosomal machinery marker, i.e. CathepsinL, p62, ATG8a (Fig. 1J,K, S1 S-X). Notably, the presynaptic accumulation phenotype could be rescued by endogenous RAB2 re-expression (Fig. 1L-N). Succeeding absence of presynaptic material from the synaptic terminals affected presynapse formation and diminished AZ number (Fig. 2A-K, S3A-G), finally resulting in impaired synaptic transmission (Fig. 2R-V). As *rab2* mutant animals were viable to the 3^rd^ instar larval stage and AZs are obviously still forming, we cannot exclude, that alternative RAB2 independent transport routes for presynaptic material might exist. RAB2 apparently functions specifically in presynaptic protein delivery, as other axonally trafficking proteins, i.e. the mitochondrial ATP-Synthase, showed neither an altered distribution at the cell body nor at the synaptic terminal (Fig. S1P-R, S3D, H). Architecture and SV number at the remaining AZs were unaffected in *rab2* mutants analyzed by STED and electron microscopy (Fig. 2H-Q), suggesting a RAB2 function in presynaptic material supply rather than incorporation into the presynapse. Of note, RAB2 function was shown to be neuron specific and cell-autonomous by neuronal RNAi knock-down (Fig. S2, S3I-R). We could further show that RAB2 associated with anterograde trafficking precursors vesicles (Fig. 3A-M) and effectively reached the synaptic terminals (Fig. 3N,O), indicating that somatic presynaptic accumulations of RAB2 mutants might be composed of precursor vesicles.

We could localize RAB2 function to the Golgi apparatus as somatic RAB2 signal overlapped with Golgi markers (Fig. 4A-C, S4P-R) and importantly, presynaptic material accumulated at the secretory *trans*-Golgi site in overlap with Syx16 (Fig. 4D-J). Notably, we observed BRP positive puncta, likely representing presynaptic precursors (Fig. 4K-M), at the Golgi, suggesting that presynaptic proteins here pass through the Golgi, likely an essential step for protein maturation through posttranslational modifications (Klumperman, 2011) as was previously suggested when studying rodent cultivated neurons, to allow for post-translational modification during protein maturation (Dresbach et al., 2006; Maas et al., 2012).

Ultrastructural analysis confirmed the vesicular nature of presynaptic material accumulation of *rab2* mutants, as we identified tubule-shaped, clear cored vesicles with a short diameter of ~40 nm accumulating in fields at the *trans*-Golgi (Fig. 5A-D). 3D tomograms showed elongated structures, which at least in partseemed to consistof smaller vesicles unable to proceed in either fusion or fission (Fig. 5G). In contrast, the previously identified presynaptic lysosome-related vesicles (PLVs) observed in *arl8* mutants (Vukoja et al., 2018) had often a more dense core and were circular with ~60 nm diameter, a 3-fold larger volume and 2-fold increased surface area (Fig. 5B-D, H,I, S5F). Collectively these findings suggest that RAB2 regulates membrane trafficking at the secretory *trans*-Golgi network to allow for early precursor formation, including both AZ scaffold protein and SV protein carrying immature PVs, a prerequisite for later maturation steps yielding mature, ~60 nm sized, circular PVs. Interestingly, we further determined by STED analysis that presynaptic proteins might sort out of the Golgi on at least two different routes as we could distinguish immature PVs with differential presynaptic cargo load in *rab2* mutants: those positive for AZ proteins and the lysosomal adaptor Arl8 and those positive for SV proteins and the lysosomal membrane protein LAMP1 (Fig. 1D-K and Fig 6D-I). Lysosomal membrane proteins traffic on non-clathrin coated vesicles, so called LMP carriers, from the TGN to the late endosome (Pols et al., 2013). It is tempting to speculate that SV and LAMP1 positive early PVs might be related to these organelles. In contrast, vesicles accumulating in *arl8* mutants had a uniform charge of BRP, VGlut and LAMP1 (Fig. 6F,G), supporting the hypothesis that PLVs of *arl8* mutants represent mature states of precursor biogenesis, while RAB2 regulates early precursor formation processes. Indeed, we could show by genetic epistasis experiments that RAB2 acts upstream of Arl8 (Fig. 7A-E). We hence suggest that presynaptic precursor biogenesis takes place in two consecutive steps: early RAB2 dependent sorting of precursor proteins from the trans-Golgi on independent pathways for AZ and SV proteins, followed by later maturation steps uniting both cargo types, either by direct fusion of early precursors or through conventional fusion into endo/lysosomal compartments, from which at a later time point mature PLVs are retrieved (Fig. 7F). An important finding represents the observation, that both the VGlut positive subtraction of early (*rab2* mutants) and the late (*arl8* mutants) presynaptic precursors are positive for the lysosomal marker LAMP1, while they are devoid of the lysosomal peptidase cathepsin, thus both share a lysosomal membrane identity, while they are clearly no degradative compartments. It remains however puzzling, that the lysosomal kinesin adaptor Arl8 localizes to the second BRP positive subfraction of early precursors as it might be expected to be component of the VGlut/LAMP1 carrying early precursors. A possible, simplistic interpretation could be that although primarily described as a regulator of lysosome and lysosome related organelle mobility (Khatter et al., 2015; Rosa-Ferreira and Munro, 2011) the elementary function of Arl8 would comprise of the directional movement of organelles with a lysosomal membrane identity, thus including, next to conventional lysosomes also mature presynaptic precursors, possibly also including SV precursor described in *C.elegans* (Klassen et al., 2010), where Arl8 was shown to affect SV precursors trafficking.

Several models exist to describe the coordinated transport of presynaptic precursors: (i) independent precursors for different cargo types, such as the 80 nm sized dense core Piccolo-bassoon transport vesicles (PTVs) for AZ proteins (Shapira et al., 2003) or STVs (Synaptic vesicle transport vesicles) with a clear core for SV proteins (Maeder et al., 2014a) or (ii) co-traffickingof AZ and SV proteins on either the same precursor or the existence of preassembled multi-vesicular transport aggregates containing dense and clear cored vesicles of unknown origin (Bury and Sabo, 2011; Tao-Cheng, 2007). We here suggest that presynaptic precursors are originally Golgi-derived vesicles requiring RAB2 as an essential regulator of membrane remodeling during this sorting process, in agreement with previous studies in mammalian cultivated neurons (Dresbach et al., 2006; Maas et al., 2012). These post-Golgi early precursors have a differential cargo load, with molecular characteristics correlating with above described PTVs or STVs, while they are two-fold smaller, not circular and with a clear core, compared the 80 nm sized dense core PTVs. Subsequent maturation steps yield mature precursors with a merged cargo load and the molecular lysosomal signature of PLVs.

The here described function of RAB2 in presynaptic precursor biogenesis could share mechanistic traits with the previously described function of RAB2 and its effectors (RIC-19, RUND-1, CCCP-1, TBC-8) in dense core vesicle (DCV) maturation in *C.elegans* (Ailion et al., 2014; Edwards et al., 2009; Hannemann et al., 2012; Sumakovic et al., 2009). DCVs are generated at the TGN (trans-Golgi network) of the Golgi (Kim et al., 2006; Morvan and Tooze, 2008), similar to the here described RAB2 dependent early precursor formation process. During the secretion process from the Golgi, constitutive, but inappropriate secretory proteins including lysosomal proteins are incorporated into the maturing DCVs and need to be removed by clathrin-mediated membrane trafficking to the endo/lysosomal system in order to finalize DCV maturation (Ailion et al., 2014; Edwards et al., 2009; Kim et al., 2006; Morvan and Tooze, 2008; Sumakovic et al., 2009). Elegant studies in *C.elegans* showed that RAB2 and RAB2 effectors are required for the retention of neuropeptides in immature DCVs during the out-sorting and removal of inadequate proteins, as in *rab2* mutants neuropeptides are miss-sorted into endo/lysosomal compartments and consequently reduced in the mature DCVs. Although we can only speculate, RAB2 might also during presynaptic precursor formation by implied into such protein sorting processes, i.e. the sorting of presynaptic and/or lysosomal proteins, although the “frozen-in-fusion-or-fission” phenotype of the Golgi-close immature precursors could as well indicate a role for RAB2 in catalyzing membrane trafficking events. However, as loss of RAB2 causes a severe presynaptic phenotype in the reduction of presynapse number and neurotransmission, which was not observed in *C.elegans*, (Edwards et al., 2009; Sumakovic et al., 2009), RAB2 function in *Drosophila* might be implied in DCV maturation, but is certainly implied in additional Golgi-related secretory processes including presynaptic precursor biogenesis.

Our findings could contribute to design a comprehensive model of the biosynthetic pathway underlying presynaptic PV biogenesis and help to unite the different models and observations of the last decades. The link of RAB2 dependent secretory Golgi function to the presynaptic TV biogenesis might help in the future to better understand RAB2 related neurodevelopmental defects, e.g. memory defects prefrontal morphology in human (Li et al., 2015), autism spectrum disorders (ASDs) or schizophrenia (SCZ) (Kiral et al., 2018; Takata et al., 2016), but also connect to Golgi-pathway related neurodegenerative diseases (Rasika et al., 2018).

## Supplemental Figures

**Fig. S1 | Presynaptic proteins accumulate in the cell bodies of *rab2*^−/−^ deficient neurons.** (A) Western Blot analysis of wild type and *rab2* mutant brains probed against RAB2 (upper panel) and alpha-tubulin (lower panel) as loading control. (B-F) Quantifications of neuronal somata comparing *rab2* mutant and wild type. Quantification of (B) RIM-BP sum intensity from Fig. 1D (wt: 100.0 ± 26.94 %, n=9; rab2 −/− : 1061 ± 241.8 %, n=9) and (C) VGlut sum intensity (wt: 100.0 ± 21.09 %, n=5; rab2 −/− : 752.0 ± 173.4 %, n=5) from Fig. 1F. (D) Quantification of BRP and VGlut aggregate area (BRP: 0.314 ± 0.02 nm, n=5; VGlut: 0.638 ± 0.07 nm; n=5) from Fig. 1F. Quantification of (E) LAMP1 (wt: 100.0 ± 39.14 %, n=5; rab2 −/− : 1322 ± 235.2 %, n=4) from Fig. 1H and (F) CathepsinL sum intensity (wt: 100.0 ± 6.05 %, n=5; rab2 −/− : 108.5 ± 14.95 %, n=5) from Fig. 1J. Confocal images of neuronal somata deficient for RAB2 stained for BRP (green) and (G) UNC13A (magenta), (J) Syt-1 (magenta), (M) Dap160 (magenta), (P) ATP-Synthase (magenta), (S) p62 (magenta), (V) ATG8a (magenta). Scalebar 2 μm. Quantification of the corresponding number of aggregates for (H) UNC13A (wt: 100.0 ± 10.66 %, n=4; rab2 −/− : 271.6 ± 25.44 %, n=4), (K) Syt-1 (wt: 100.0 ± 2.52 %, n=4; rab2 −/− : 161.4 ± 16.69 %, n=4), (N) Dap160 (wt: 100.0 ± 21.24 %, n=4; rab2 −/− : 1927 ± 183.1 %, n=5), (Q) ATP-Synthase (wt: 100.0 ± 12.38 %, n=4; rab2 −/− : 107.2 ± 16.89 %, n=4), (T) p62 (wt: 100.0 ± 11.43 %, n=5; rab2 −/− : 81.92 ± 7.939 %, n=5) and (W) ATG8a (wt: 100.0 ± 15.09 %, n=4; rab2 −/− : 89.01 ± 9.307 %, n=4). Quantification of the corresponding protein sum intensity for (I) UNC13A (wt: 100.0 ± 19.18 %, n=4; rab2 −/− : 319.7 ± 39.60 %, n=4), (L) Syt-1 (wt: 100.0 ± 14.46 %, n=4; rab2 −/− : 176.5 ± 14.63 %, n=4), (O) Dap160 (wt: 100 ± 22.5 %, n=4; rab2 −/− : 10300 ± 1360 %, n=5), (R) ATP-Synthase (wt: 100.0 ± 14.14 %, n=4; rab2 −/− : 94.48 ± 10.85 %, n=4), (U) p62 (wt: 100.0 ± 10.37 %, n=5; rab2 −/− : 86.99 ± 17.72 %, n=5) and (X) ATG8a (wt: 100.0 ± 25.09 %, n=4; rab2 −/− : 93.91 ± 13.78 %, n=4). All graphs show mean +/− SEM. Normality was tested (for data sets with n>4) with the Kolmogorov-Smirnov test, if normal distributed (or assumed to be normally distribute for n<5) unpaired t-test was used (B-E, H+I, K+L, N+O, Q+R, T+U, W+X), otherwise non-parametric Mann-Whitney test (F). *p<0.05, **p<0.01, ***p<0.001.

**Fig. S2| RAB2 operates neuron specific in a cell autonomous manner.** (A) Western Blot analysis of wild type and Rab2 RNAi knock down brains expressing the RNAi specifically in motoneurons (ok6-Gal4 driver) probed against RAB2 (upper panel) and alpha-tubulin (lower panel) as loading control. (B) Confocal images of neuronal somata from control(driver control) and Rab2 RNAi knock down brains immunostained for the AZ scaffold protein BRP (green). Neuronal cell bodies in the VNC cortex (dotted lines). White squares show zoom area. Scale bar overview 10 μm, zoom 2 μm. (B,C) Quantifications of the representative images of (B). (C) Number of BRP aggregates (ctrl: 100.0 ± 8.785 %, n=3; Rab2-RNAi: 180.1 ± 24.43 %, n=3). (D) BRP sum intensity (ctrl: 100.0 ± 18.38 %, n=3; Rab2-RNAi: 346.6 ± 38.85 %, n=3). (E) Confocal images of neuronal somata of control (driver control) and Rab2 RNAi knock down brains immunostained for BRP (green) and VGlut (magenta). (F) Quantifications of number of VGlut aggregates (ctrl: 100.0 ± 16.85 %, n=4; Rab2-RNAi: 156.9 ± 11.86 %, n=4) of the representative images of (E). (G) Confocal images of neuronal somata of control (driver control) and Rab2 RNAi knock down brains immunostained for BRP (green) and Syt-1 (magenta). (H) Quantifications of number of Syt-1 aggregates (ctrl: 100.0 ± 33.50 %, n=3; Rab2-RNAi: 367.6 ± 80.73 %, n=3) of the representative images of (G). Scale bar for (E) and (G) 2 μm. All graphs show mean +/− SEM. Normal distribution was assumed but not formally tested for n<5 (C,D,F,H). *p<0.05, **p<0.01, ***p<0.001.

**Fig. S3| Presynaptic biogenesis relies on the RAB2 dependent delivery of presynaptic material.** Confocal images of wild type and *rab2* mutant NMJs stained in green for (A) UNC13A, (B) Syt-1, (C) Dap160, (Di) ATP-Synthase and (Dii) ATP-Synthase (green) and HRP (magenta, as a neuronal membrane marker). Scalebar 3 μm. Corresponding quantification of the protein sum (or mean pixel) intensity for (E) UNC13A (wt: 100.0 ± 15.83 %, n=10; rab2 −/− : 28.96 ± 4.821 %, n=10), (F) Syt-1 (wt: 100.0 ± 19.60 %, n=8; rab2 −/− : 53.57 ± 4.89 %, n=8), (G) Dap160 (wt: 100.0 ± 20.83 %, n=8; rab2 −/− : 19.35 ± 2.592 %, n=10) and (H) ATP-Synthase (wt: 100.0 ± 5.060 %, n=8; rab2 −/− : 129.4 ± 15.54 %, n=7). (I) Confocal images of NMJs from control (driver control) and Rab2 RNAi knock down terminals immunostained for the AZ scaffold protein BRP(green) and HRP(magenta). (J) corresponding zoom images. Scale bar overview 5 μm, zoom 2 μm. (K-M) Quantifications of the representative images of (I). (K) BRP sum intensity (ctrl: 100.0 ± 5.74 %, n=7; Rab2-RNAi: 54.92 ± 4.04 %, n=7). (L) Number of AZ (ctrl: 100.0 ± 6.0 %, n=7; Rab2-RNAi: 83.43 ± 3.772 %, n=8) and (M) AZ area (ctrl: 100.0 ± 6.436 %, n=7; Rab2-RNAi: 101.2 ± 3.556 %, n=8). (N-O) Two electrode voltage-clamp electrophysiological recordings of NMJs from control (driver control) and Rab2 RNAi knock down larvae. (N) Representative example eEJC traces of control and RAB2 knock-down synapses. (O) eEJC amplitudes (ctrl: −67.18 ± 4.48 nA, n=10; Rab2-RNAi: −42.61 ± 3.408 nA %, n=10). (P) eEJC charge (ctrl: −642.7 ± 48.07 pC, n=10; Rab2-RNAi: −393.9 ± 27.95 pC, n=10). (Q) eEJC tau (ctrl: 5.94 ± 0.24 ms, n=10; Rab2-RNAi: 5.65 ± 0.17 ms, n=10). (R) 10 ms paired-pulse ratio (ctrl: 1.01 ± 0.11, n=10; Rab2-RNAi: 1.01 ± 0.13, n=10). All graphs show mean +/− SEM. Normality was tested with the Kolmogorov-Smirnov test, if normal distributed unpaired t-test (E, G, H, K, L, M, O-Q) was used, otherwise non-parametric Mann-Whitneytest (F, R). *p<0.05, **p<0.01, ***p<0.001.

**Fig. S4| RAB2 localizes to the Golgi in neuronal somata.** Confocal images of neuronal somata of larvae expressing RAB2-YFP (green) into the motoneurons co-stained in magenta for (A) Rbsn-5, (D) Rab7, (G) Dor, (J) CathepsinL, (M) ATG8a and (P) Syntaxin 16 (Syx16). Scale bar 5 μm. Corresponding line profiles for (B) Rbsn-5, (E) Rab7, (H) Dor, (K) CathepsinL, (N) ATG8a and (Q) Syx16. Corresponding Pearson’s correlation coefficients for (C) Rbsn-5 (Rab2-YFP vs. Rbsn-5: −0.26 ± 0.04, n=11; inverted Rbsn-5: −0.23 ± 0.04, n=11), (F) Rab7 (Rab2-YFP vs. Rab7: −0.23 ± 0.05, n=12; inverted Rab7: −0.13 ± 0.053, n=12), (I) Dor (Rab2-YFP vs. Dor: −0.2 ± 0.04, n=12; inverted DOR: −0.13 ± 0.04, n=12), (L) CathepsinL Rab2-YFP vs. CathepsinL: −0.14 ± 0.05, n=10; inverted CathepsinL: −0.13 ± 0.07, n=10), (O) ATG8a (Rab2-YFP vs. ATG8a: −0.21 ± 0.04, n=12; inverted ATG8a: −0.24 ± 0.03, n=12). (R) Quantification of the area overlap of Rab2-YFP and Syx16 in % (Rab2 and Syx16: 88.38 ± 2.68, n=9; inverted Rab2 and Syx16: 7.91 ± 2.36, n=9), n represents single neuronal somata from 3 animals/brains, 3 cells/brain. Allgraphs show mean +/− SEM. Normality was tested with the Kolmogorov-Smirnov test, if normal distributed unpaired t-test (C, F, L, O, R) was used, otherwise non-parametric Mann-Whitney test (I). *p<0.05, **p<0.01, ***<0.001.

**Fig. S5| Tubule-shaped vesicular membranes accumulate at the *trans*-Golgi in *rab2* mutants.** (A,B coloured) Electron micrographs of wild type and *rab2* mutant whole cell bodies containing Golgi(Go) areas (N = nucleus) with vesicle accumulations at the *trans*-Golgi(green) in *rab2* deficient neurons. Scale bar 500 nm. (C) Quantification of long diameter of accumulating vesicles in *rab2* (Fig. 5B middle panel) and *arl8* (Fig. 5B right panel) mutants (rab2−/−: 72.07 nm ± 1.27, n= 313; arl8−/−: 78.70 nm ± 1.54, n=138). N represents single vesicles. (D) Volume fraction of autophagocytic vacuoles in wild type and RAB2 deficient somata (wild type: 0.0007 ± 0.0003,n= 39; rab2 −/−: 0.0021 ± 0.0005,n=73). (E) Virtual section and 3D tomography of a wild type neuronal somata (left) and 3D tomography reconstruction (right) (colours). Scale bar 0.2 μm. (F) Surface area of accumulating vesicles of *rab2* and *arl8* mutant neuronal cell bodies (rab2−/−: 33563 nm^2^ ± 794, n= 313; arl8−/−: 60795 nm^2^ ± 1508, n=138). N represents single vesicles.

**Fig. S6| LAMP1 positive precursors localize at the Golgi and precursor independent LAMP1 accumulations in *rab2* mutant brains.** STED imaging of a representative PV accumulations in *rab2* mutant somata immunostained for GFP::LAMP1 (magenta) and (A) GM130 (green) or (C) BRP (green). Scale bar 0.5 μm. (B) Line profile of (A). (D) Quantification of VNC aggregates positive for LAMP1 only and co-positive for LAMP1 and BRP in % (LAMP1 only positive aggregates: 46.01 % ± 4.35, n=25; LAMP1 and BRP positive aggregates: 53.99 % ± 4.35, n=28). N represents single aggregates from 3 larvae and 4 −5 VNC images/animal.

## Materials and Methods

### Contact for Reagent and Resource Sharing

Further information and requests for resources and reagents should be directed to and will be fulfilled by the Lead Contact Astrid Petzoldt.

### Experimental Model and Subject Details

#### Drosophila melanogaster

Fly strains were reared under standard laboratory conditions and raised at 25°C and 70% humidity on semi-defined medium (Bloomington recipe). For RNAi experiments flies were kept at 29°C. For electrophysiological recordings, only male larvae were used, for all other experiments both male and female animals were used. See Key Resources Table for genotypes and strains used.

Sample size estimation: No estimation of simple size was done as sample sizes were not chosen based on pre-specified effect size. Instead, multiple independent experiments were carried out using several biological replicates specified in the legends to figures.

## Method Details

### Immunostainings of larval NMJs and brains

#### Immunohistochemistry for confocal and STED microscopy

For immunohistochemistry dissections were performed in haemolymph-like solution 3 (HL3; (Stewart et al., 1994); composition in mM: 70 NaCl, 5 KCl, 20 MgCl_2_, 10 NaHCO_3_, 5 trehalose, 115 sucrose, 5 HEPES, pH adjusted to 7.2) by opening the 3^rd^ instar larvae dorsally along the midline and removing the innards. Dissections were fixated with 4 % para formaldehydein PBS (pH 7.2) for NMJ stainings or Bouin’s fixative for brain stainings for 10 min. After fixation, the filets were washed with PBS plus 0.05 % Triton-X 100 (PBT) and blocked for 60 min in 5% normal goat serum (NGS). For immunostainings, the larvae were incubated with primary antibodies at 4° C overnight and subsequently washed in a 0.05 % PBT solution for 2 h at room temperature (RT). Larvae were then incubated for 2-3h with secondary antibodies at RT. Washing procedures were repeated. Larvae were finally mounted either in Vectashield (Vector Laboratories), Mowiol (SigmarAldrich) or ProLong Gold (Thermofischer).

#### Image acquisition and analysis

Conventional confocal and STED images were acquired with Leica DMI 6000 (SP8) and TCS SP8 gSTED 3× microscopes (Leica Microsystems), respectively. For confocal scans a HC PL APO CS2 63x /1.40-N.A. oil objective (Leica Microsystems) was used, for STED a HC PL APO CS2 100×/1.40-N.A. oil objective (Leica Microsystems). Images were acquired at approximately 20°C and used fluorochromes are indicated in the antibody section.For detection HYD (high sensitive) 400-800 nm spectralde scanned for green and red channels and PMT 400-800 nm spectral descanned for far red channels were used for confocal scans. For STED HyD Sp GaAsP were used. Larval brain z-stacks had a step size of 0.3-0.5 μm between single optical slices. 40 z-slices of the central region along the dorso-ventral axis were imaged. The NMJ z-stacks had a step size of 0.2 – 0.3 μm between single optical slices. All images were acquired using the LAS X software (Leica Microsystems, Wetzlar, Germany). For all confoca land STED image analysis the software ImageJ 1.52n was used, for all statistical analysis the software GraphPad PRISM, version 5.01. For STED microscopy Huygens Deconvolution software was used applying a theoretical point spread function automatically computed based on pulsed- or continuous-wave STED optimized function and the specific microscope parameters. Default deconvolution settings were applied. Images for figures were processed if necessary, with ImageJ software to enhance brightness using the brightness/contrast function and smoothened (0.5 pixel Sigma radius) using the Gaussian blur function. Confocal stacks were processed with Fiji (http://fiji.sc) (Schindelin et al., 2012). Image analysis followed the standard protocol as described in (Andlauer and Sigrist, 2012b) and will be described in detail for the different analyses as follows.

For NMJ quantification the signal of the HRP-Cy5 antibody was used as template for a mask, restricting the quantified area to the shape of the NMJ. For larval brain aggregate quantification, a ROI (region of interest) was drawn via the freehand selection tool including only the cortex region of the larval ventral nerve cord (sparing neuropil region and background). The original confocal stacks were converted to maximal projections (for brain taking only the central 20 z slices of the ventral nerve cord without adjacent axons or weakly stained tissue in deeper layers), and after background subtraction, a mask of the synaptic area / brain aggregates was created by applying a certain threshold to remove the irrelevant lower intensity pixels. The segmentation of single spots was done semi-automatically via the command “Find Maxima” embedded in the Fiji and by hand with the pencil tool and a line thickness of 1 pixel. To remove high-frequency noise a Gaussian blur filter (0.5 pixel Sigma radius) was applied. The processed picture was then transformed into a binary mask using the same lower threshold value as in the first step. This binary mask was then projected onto the original unmodified image using the “min” operation from the ImageJ image calculator. For sum intensities all intensities of the corresponding channel of one NMJ / larval brain were added up, n represents the number of NMJs / larval brains.

Pearson’s correlation coefficient image stack was acquired, and one focal plane chosen for analysis after ROI selection of a single cell. ImageJ Coloc_2 was used to determine the Pearson’s R value above threshold. As control one channel was flipped horizontally to show random co-localization of the image. Analyzed where overall 9-12 cells of 2-3 VNCs. BRP ring diameter (C-terminal antibody Nc82) from STED analysis of NMJs: Nc82 positive active zones in a planar position with a minimum of three clusters/maxima were manually encircled and ROIs were saved (1-14 AZs/bouton, 3 boutons/larvae, 3 larvae). ROI area was measured (Fiji) and diameter calculated assume a circular ROI area according to the formula: d=2 * square root (A/π), with d=diameter, A=area.

BRP mean pixel intensity in boutons excluding AZs from STED analysis of NMJs: Bouton was outlined manually (ROI-1) and outside bouton area was cleared. Gaussian blur with 2.0 was applied. A threshold of 30 was set and all AZ within this threshold marked (ROI-2). Subtraction of ROI-2 from ROI-1 (XOR function in Fiji ROI manager) and in the new ROI-3 (bouton area without AZs) mean pixel intensity measured. Data from 3 boutons/larvae, 3 larvae.

Number of AZper bouton area from STED analysis of NMJs: ROI-1 (see above) was measured and area determined. All AZs (planar and lateral, with a minimum of two clusters/maxima) were counted manually, data from 3 boutons/larvae, 3 larvae.

For area overlap of Rab2-YFP and Syx16 in % from neuronal somata, single cells were manually chosen (3/brain, 3 brains in total) and a Gaussian Blur of 1.0 was applied. Threshold values were set manually and ROIs for in both channels marked and area measured. The AND function of the Fiji ROI manager was used to determine area overlap. Rab2-YFP image was flipped vertically or horizontally and area overlap determined as above.

Frequency in % of peak to peak distance of BRP positive aggregates in *rab2* mutants and GM130 marked Golgi: place line through BRP aggregates and Golgi, save as ROI, use ROI manager MultiPlot tool and determine peak to peak distance. For random shifted data, BRP channel was shifted by 80 pixels. Cut off at 1.2 μm distance, binned for 0.06 μm. With 1 −14 measured distances from 2 VNC images/larvae, with 3 larvae total analyzed.

Center distance of STED imaged presynaptic protein aggregates in *rab2* mutant somata: Single channel images were created for individual aggregates. By defining a suitable threshold for each aggregate separately, a binary mask was created, outlines of the aggregate were detected by the magical wand tool and subsequently saved as ROI. Using the ImageJ built-in measurement function “center of mass” the respective x and y mass-central coordinates of each ROI (one per channel) were calculated. Distance of the centers (centers of mass) was calculated according to the function d=squareroot((x(channel1)-x(channel2)^2+(y(channel1)-y(channel2))^2). N represents the number of single aggregates tested in ventral nerve cord images of 2-3 animal.

### *In vivo* live imaging and analysis of *Drosophila* larvae

*In vivo* imaging of intact *Drosophila* larvae was performed as previously described (Andlauer and Sigrist, 2012a). Briefly, third instar larvae were put into a drop of Voltalef H10S oil (Arkema, France) within an airtight imaging chamber. The larvae were anaesthetized before imaging with 10 short pulses of a desflurane (Baxter,IL, UAS) air mixture until the muscles relaxed. Axons immediately after exiting the ventral nerve cord were imaged using confocal microscopy. Kymographs were plotted using a custom-written Fiji script (see STAR method table).

### Electrophysiology

Two-electrode voltage clamp (TEVC) recordings were carried out essentially as previously reported (Matkovic et al., 2013). They comprised spontaneous recordings (miniature excitatory junction currents, mEJC, 90 s), single evoked (evoked excitatory junction currents, eEJC, 20 repetitions at 0.2 Hz) and high-frequency recordings (paired pulse 10 ms or 30 ms interstimulus interval, PP10 or PP30, 10 repetitions at 0.2 Hz). All experiments were performed on third instar larvae raised at 25°C on semi-defined medium (Bloomington recipe). Dissection and recording medium was extracellular haemolymph-like solution 3 (HL3; (Stewart et al., 1994); composition in mM: 70 NaCl, 5 KCl, 20 MgCl_2_, 10 NaHCO_3_, 5 trehalose, 115 sucrose, 5 HEPES, pH adjusted to 7.2). Dissection was done in ice cold Ca^2+^-free HL3 medium while mEJC, eEJC and high-frequency recordings were performed in 1.5 mM Ca^2+^ HL3 at room temperature. For all physiological recordings intracellular electrodes with a resistances of 15−25 MΩ (filled with 3M KCL) were placed at muscle 6 of abdominal segment A2 / A3. Acquired data was low-pass filtered at 1 kHz and sampled at 10 kHz. Traces of both miniature and evoked postsynaptic currents were recorded in TEVC mode (AxoClamp 2B; Axon Instruments, Union City, CA). The command potential (V_cmd_) for mEJC recordings was −80 mV and −60 mV for all other recordings. Only cells with an initial membrane potential (*V*_m_) between −50 and −70 mV and input resistances of ⩾4 MΩ were used for further analysis. eEJC and paired-pulse traces were analyzed for standard parameters (amplitude, rise time, decay, charge flow, PP-ratio) by using a semi-automatic custom-written Matlab script (Mathworks, vers. R2009a). Stimulation artifacts in eEJC and paired-pulse recordings were removed for clarity. mEJC recordings were analysed with pClamp 10 software (Molecular Devices).

### Western blot analysis of larval brains

Larval CNS protein extraction were performed as follows: 10 CNS were dissected from 3rd instar larvae. The obtained tissues were sheared manually in 5 μl of 2% SDS aqueous solution using a micropistil fitting tightly into a 1.5-ml cup. An amount of 0,5 μl of a 10% Triton-X 100 aqueous solution and 5 μl of 2× Laemmli sample buffer was added, and samples were heated at 95°C for 10 min. After centrifugation for 2 min at 16,000×g, in order to pellet the debris. For larval CNS 10 μl (equivalent to 10 larval CNS) was subjected to denaturing SDS-PAGE using an 6% Tris_HCl gels.

Proteins were then transferred onto a nitrocellulose membrane blocked with 5% skim-milk in 1× PBS supplemented with 0.1% Tween-20 and probed with rabbit anti-Rab2 (1:5000) (Santa Cruz Biotechnology, sc-285667) and mouse anti-Tubulin (1: 100000) diluted in 5% milk in 1x PBS, supplemented with 0.1% Tween-20. After washing, secondary anti-rabbit or anti-mouse IgG horseradish peroxidase (HRP)-conjugated antibodies (Dianova) were used for detection (Dianova) in conjunction with an enhanced chemoluminescence (GE Healthcare ECL Prime; product number RPN 2232) detection system with Hyperfilm ECL (GE Healthcare).

### Electron microscopy

Conventional embedding was performed as described previously (Matkovic et al., 2013). In brief, dissected third instar larvae were fixed with PFA (10 min; 4% PFA and 0.5% Glutaraldehydein 0.1 MPBS) and Glutaraldehyde(60 min; 2% glutaraldehyde in 0.1 M sodium cacodylate), washed in sodium cacodylate buffe and postfixed with 1% osmium tetroxide and 0.8% KFeCn in 0.1 M sodium cacodylate buffer (1h on ice). After washing with sodium cacodylate buffer and distilled water, the samples were stained with 1% uranyl acetate in distilled water. Samples were dehydrated, infiltrated in Epon resin and subsequently muscle 6/7 of abdominal segment A2/3 were cut out. Collected in an embedding mold, blocks were polymerized and cut in 65-70 nm thin serial sections. These sections were postfixed and poststained with uranyl acetate / lead citrate. Micrographs were taken with an electron microscope (JEM 1011; JEOL) equipped with a camera (Orius 1200A; Gatan) using the DigitalMicrograph software package (Gatan).

For quantification the plasma membrane and the electron-dense T-bar was detected by eye and labeled manually. T-bar roof size was measured by a straight line connecting the furthest distance of the upmost T-bar dense material (in relation to the plasma membrane). T-bar area was obtained by surrounding the dense material and measuring the area of the created ROI. For 2D morphometric analysis images of neuronal soma from VNC where imaged at Zeiss 900 TEM. For 3D analysis, 250 nm sections were cut and collected on coated slotted grids with 10 nm gold fiducials. Once Golgi was located, series of images from +60º to −60º were taken with a 1º step at Tecnai G20 microscope. Etomo/IMOD (Kremer et al., 1996) (https://bio3d.colorado.edu/imod/) and Microscopy imaging browser MIB (Belevich et al., 2016) (http://mib.helsinki.fi/index.html) have been used to work with 3D volume and render 3D models of Golgi and Golgi associated structures.

### Quantification and Statistical Analysis

Data were analyzed using GraphPad Prism v5.01. To compare two groups, two-tailed *t*-test (data normally distributed) or Mann-Whitney *U* test (data not normally distributed) was used. To compare three or more groups one-way ANOVA (+ Tukey’s post hoc test; data normally distributed) or the Kruskal-Wallis Test (+ Dunn’s post hoc test; data not normally distributed) was used. Data distribution was tested by the Kolmogorov Smirnov test for a sample size of n > 4. For a sample size of n < 5 normal distribution was assumed. No statistical methods were used to predetermine sample sizes, but our sample sizes are similar to those generally employed in the field. Data collection and analyses were not performed blind to the conditions of the experiments, nor was data collection randomized. For electrophysiological recordings, genotypes were measured in an alternating fashion on the same day and strictly analyzed in an unbiased manner.

## Acknowledgements

We thank Veronika Lysiuk for help with the 3D EM tomography analysis. We thank Marta Maglione and Niclas Gimber for valuable discussions and support. I thank Sebastian Schütte for support and encouragement. TWBG was financed by SFB 740, C09 und SFB 1315, A08. A.G.P. was supported by the SFB958 and the BIH gender equality fund.

## Author contributions

T.W.B.G., A.G.P. performed immunofluorescence and genetic experiments, including analysis. A.G.P. performed STED imaging and analysis. T.W.B.G. performed electrophysiological experiments. J.L. performed Western Blot analysis. D.P., C.Q. performed electron microscopy experiments, D.P. and A.G.N performed electron microcopy analysis. M.L. contributed to STED imaging and analysis. S.J.S. and A.G.P designed the study and wrote the manuscript.

