## Supplements for "Presynaptic precursor vesicles originate from the *trans*-Golgi network, promoted by the small GTPase RAB2"

**Fig. S6| LAMP1 positive precursors localize at the Golgi and precursor independent LAMP1 accumulations in *rab2* mutant brains.** STED imaging of a representative PV accumulations in *rab2* mutant somata immunostained for GFP::LAMP1 (magenta) and (A) GM130 (green) or (C) BRP (green). Scale bar 0.5 µm. (B) Line profile of (A). (D) Quantification of VNC aggregates positive for LAMP1 only and co-positive for LAMP1 and BRP in % (LAMP1 only positive aggregates: 46.01 % ± 4.35, n=25; LAMP1 and BRP positive aggregates: 53.99 % ± 4.35, n=28). N represents single aggregates from 3 larvae and 4-5 VNC images/animal.

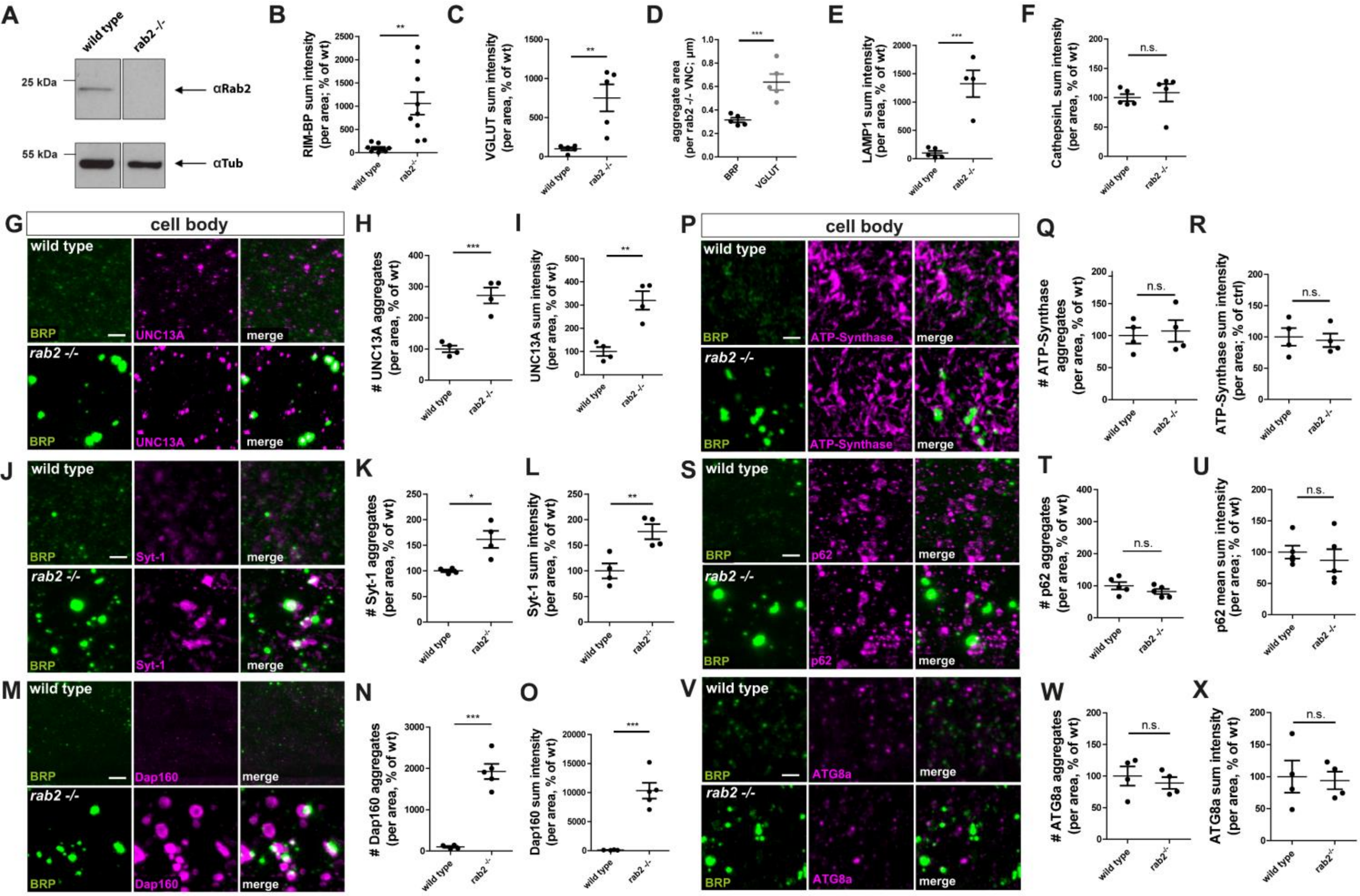


**Götz et al. , Fig. S1**

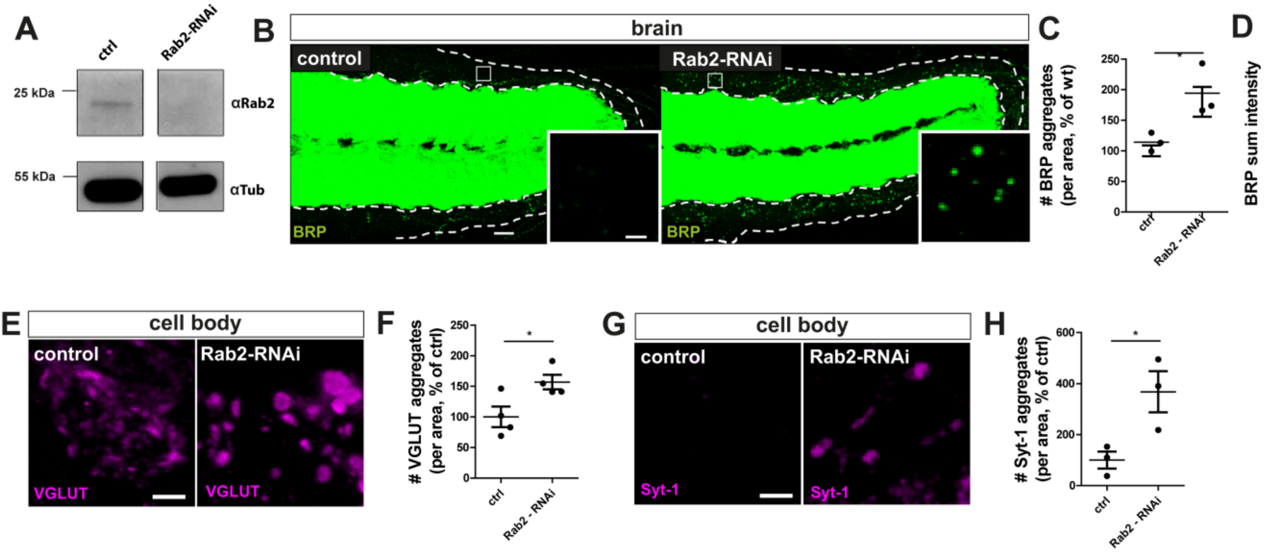


**Götz et al. , Fig. S2**

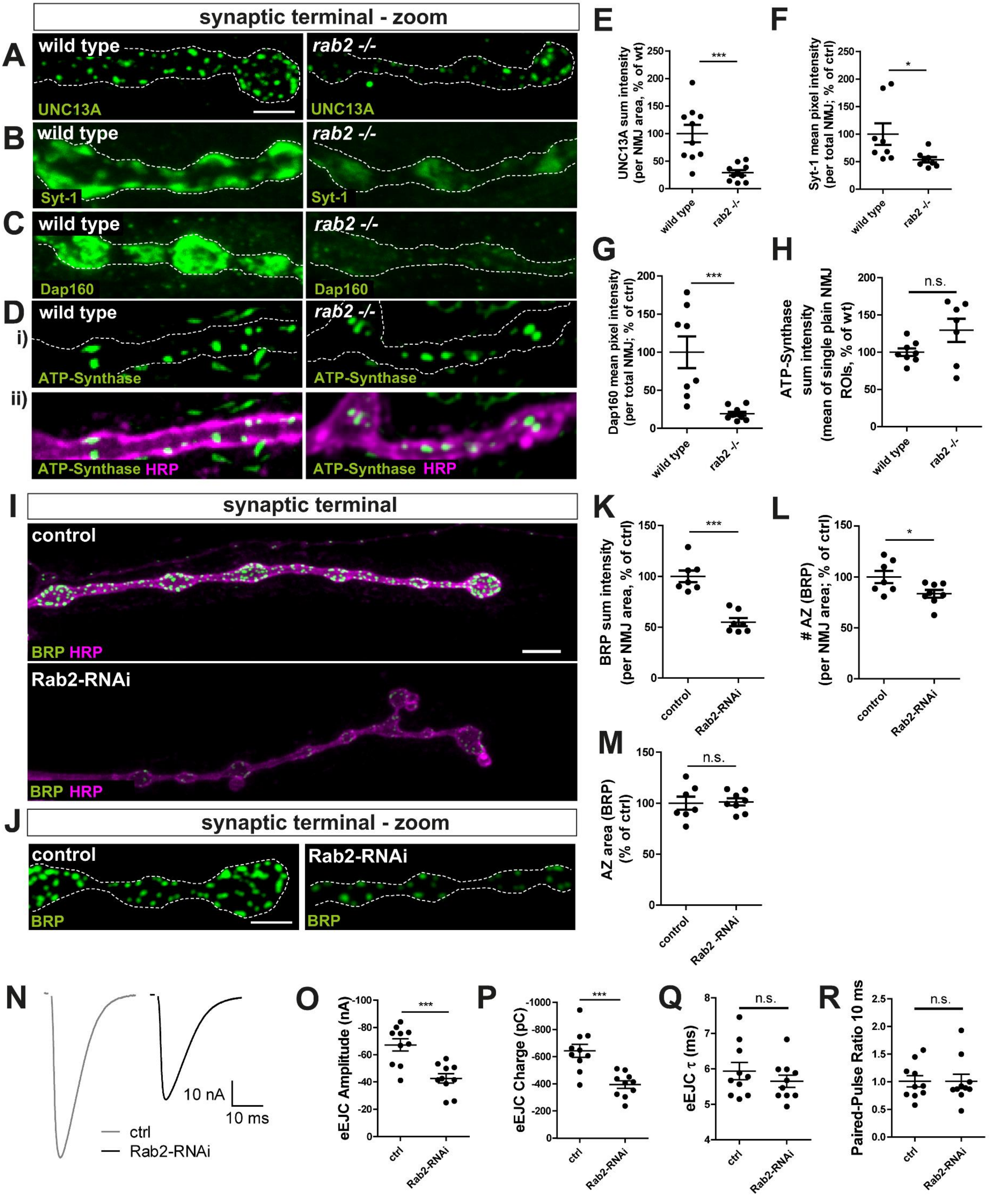


Götz et al. , Fig. S3

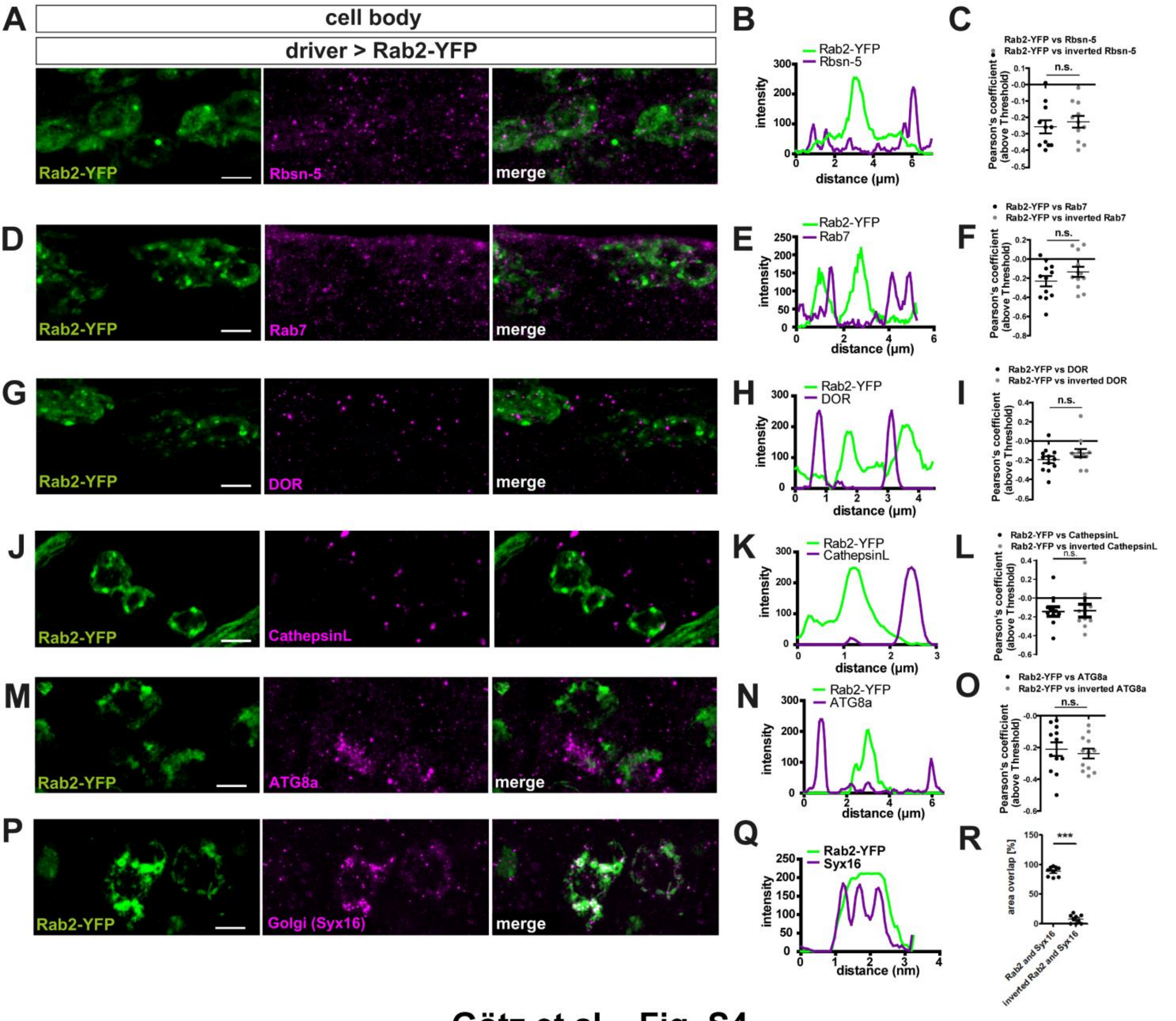


Götz et al. , Fig. S4

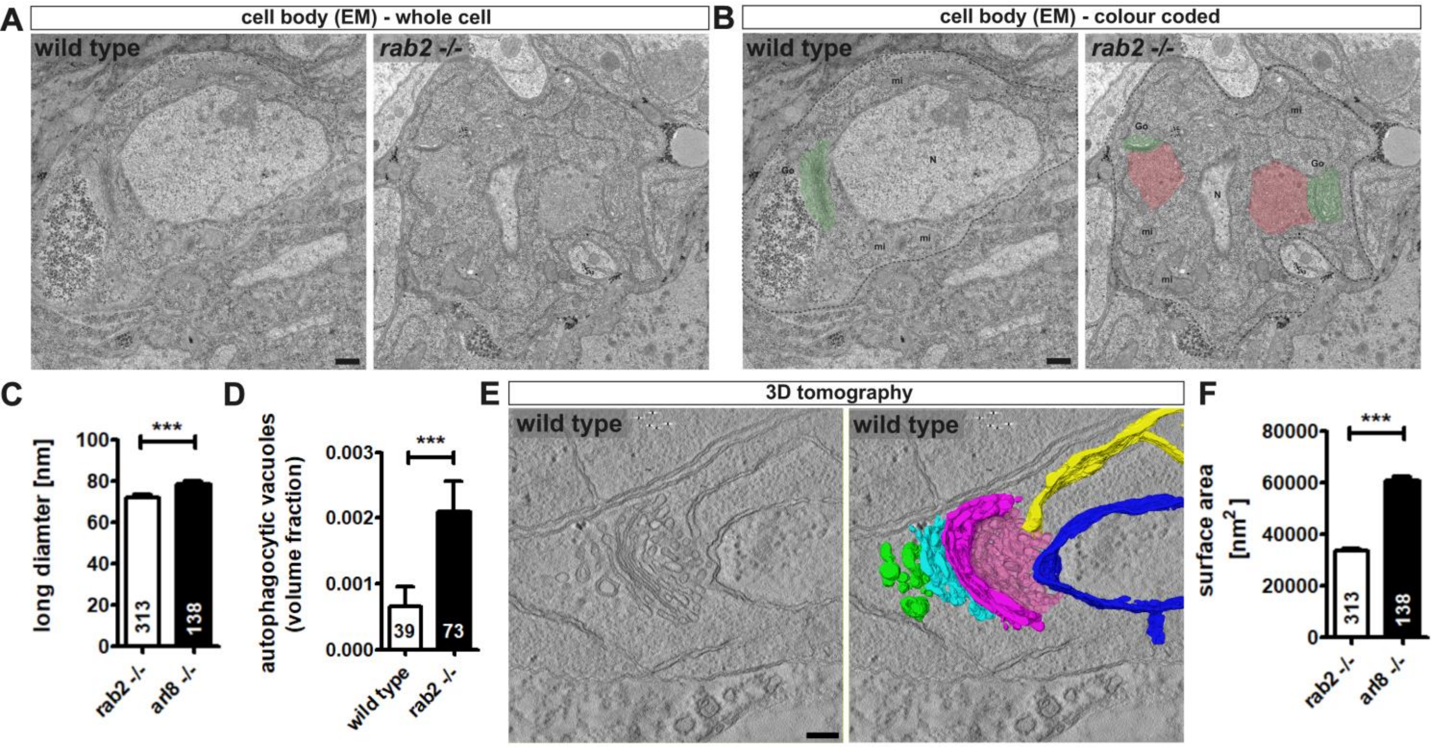


Götz et al. , Fig. S5

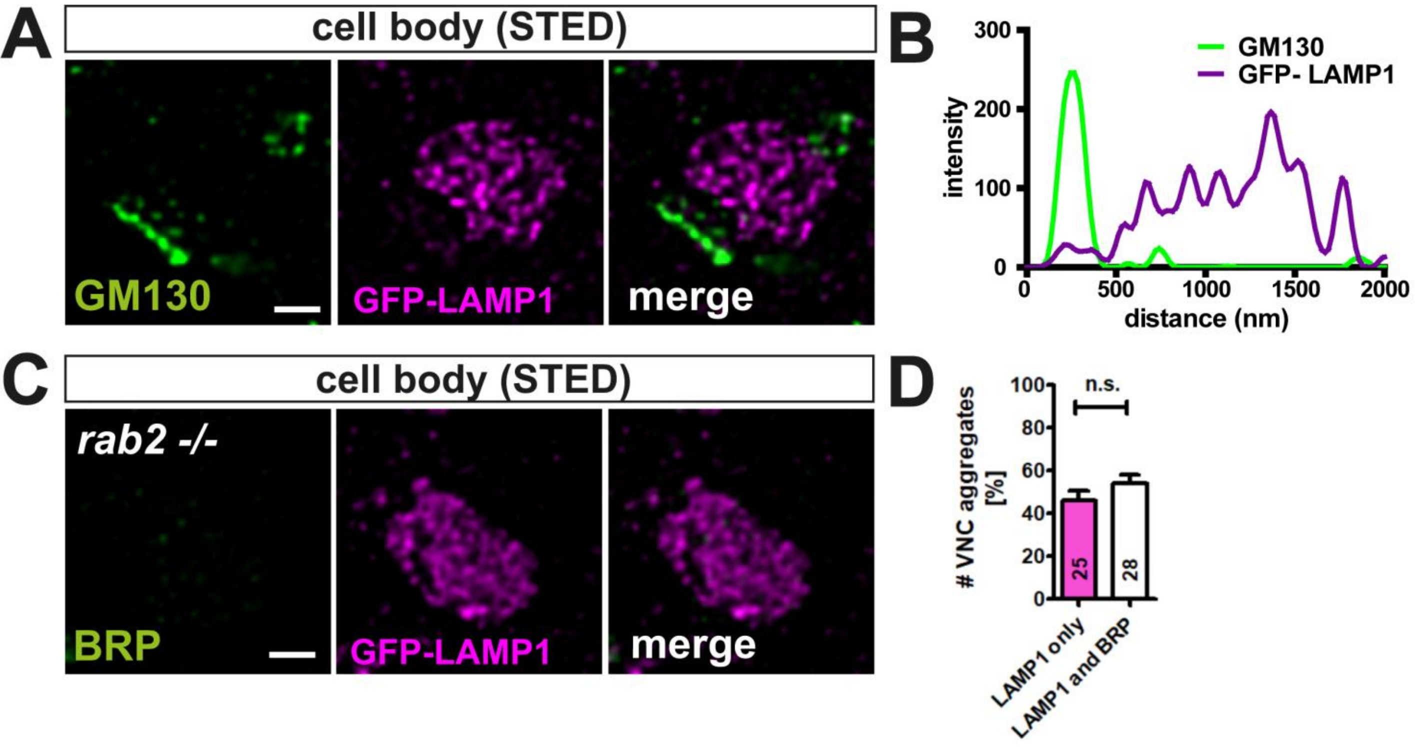


### Götz et al. , Fig. S6
